# Accurate cryo-EM protein particle picking by integrating the foundational AI image segmentation model and specialized U-Net

**DOI:** 10.1101/2023.10.02.560572

**Authors:** Rajan Gyawali, Ashwin Dhakal, Liguo Wang, Jianlin Cheng

**Affiliations:** Department of Electrical Engineering and Computer Science, University of Missouri, Columbia, MO, 65211, USA; NextGen Precision Health, University of Missouri, Columbia, MO, 65211, USA; Laboratory for BioMolecular Structure (LBMS), Brookhaven National Laboratory, Upton, NY, 11973, USA

## Abstract

Picking protein particles in cryo-electron microscopy (cryo-EM) micrographs is a crucial step in the cryo-EM-based structure determination. However, existing methods trained on a limited amount of cryo-EM data still cannot accurately pick protein particles from noisy cryo-EM images. The general foundational artificial intelligence (AI)-based image segmentation model such as Meta’s Segment Anything Model (SAM) cannot segment protein particles well because their training data do not include cryo-EM images. Here, we present a novel approach (CryoSegNet) of integrating an attention-gated U-shape network (U-Net) specially designed and trained for cryo-EM particle picking and the SAM. The U-Net is first trained on a large cryo-EM image dataset and then used to generate input from original cryo-EM images for SAM to make particle pickings. CryoSegNet shows both high precision and recall in segmenting protein particles from cryo-EM micrographs, irrespective of protein type, shape, and size. On several independent datasets of various protein types, CryoSegNet outperforms two top machine learning particle pickers crYOLO and Topaz as well as SAM itself. The average resolution of density maps reconstructed from the particles picked by CryoSegNet is 3.32 Å, 7% better than 3.57 Å of Topaz and 14% better than 3.85 Å of crYOLO.

## Introduction

Protein structure determination is a significant area of research in the field of structural biology and bioinformatics, enabling researchers to understand the roles of proteins in various biological processes^1^. This structural insight is important for studying the interaction of proteins with other molecules in the cellular processes. It is useful for finding the potential binding sites for drug molecules to act on to modulate the function of proteins^2,3^. Further, many diseases are the result of protein misfolding and aggregation. Thus, it is imperative to determine the protein structure for understanding protein function and interaction, studying their roles in the diseases, and accelerating the design of drugs.

X-ray crystallography, nuclear magnetic resonance (NMR), and cryo-EM^4,5^ are three main experimental techniques to determine protein structures. Among them, cryo-EM is the cutting-edge technique for solving the structure of large protein complexes. With advancements in electron microscope and detector devices, cryo-EM has revolutionized the field of structural biology and enabled the determination of very large protein complex structures at near atomic resolution that other experimental techniques cannot handle.

The cryo-EM-based structure determination process^6,7^ involves sample preparation with vitreous ice, imaging them with electron dose from the microscope to generate 2D projections of the samples at different orientations, followed by protein particle picking in cryo-EM micrographs (images). Once the particles are picked and extracted, the single particle analysis is employed to determine the 3D structure of the specimen.

Particle picking in cryo-EM micrographs has posed significant challenges due to the low contrast of micrographs with a low signal to noise ratio (SNR) caused by using limited electron dose during imaging process. Further, the prevalence of ice contamination, carbon edges, protein aggregates and deformed particles have further complicated the particle picking. Reconstructing a 3D protein structure from cryo-EM micrographs requires thousands of extracted particles of good quality, and therefore it is important to pick protein particles accurately and automatically, releasing the burden of human intervention and reducing the bias and inconsistency associated with manual particle picking.

With advancements in hardware and software tools^8–12^, numerous semi-automated or automated approaches varying from traditional computational methods to modern deep learning techniques have been proposed to streamline the cryo-EM processing and particle picking. Conventional computer vision methods like edge detection, blob detection and template matching^4^ are still widely used for particle picking. However, due to the low SNR of cryo-EM micrographs, these techniques are susceptible to picking ice patches, carbon areas and aggregated particles, resulting in a high number of false positives. RELION^11^ leverages a regularized likelihood optimization technique and utilizes the template-based and blob-based picking^13^ approaches. In the template-based approach, an initial set of 2D templates are generated from the manually picked particles, which are used to correlate with the different regions of micrographs to extract similar patches. This approach is highly sensitive to noise and may introduce significant bias. Similarly, in the blob-based picking, the regions of high intensity and local maxima are extracted from cryo-EM micrographs using Laplacian of Gaussian. This method is useful if the particles have significant contrast difference with the background of the micrographs and all the particles within the micrograph are of similar shape and size. If the particles are of different conformations and size, this method faces a lot of difficulty in picking the true protein particles. Other conventional tools like EMAN2^10^, SPIDER^14^, XMIPP^15^ utilizing similar computer vision approaches require a lot of manual intervention, computational resources, memory, and human time and face significant challenges of filtering out false positives.

Recent advancements in machine learning, particularly deep learning, have shown great potential for particle picking. Several machine learning approaches have been put forth to automate the particle picking process and reduce the number of false positives. Notable approaches include APPLE picker^16^, crYOLO^17^, PIXER^18^, WARP^19^, Topaz^20^, CASSPER^21^, AutoCryoPicker^22^, DeepCryoPicker^23^, DRPnet^24^ and CryoTransformer^25^. They utilize either convolutional neural networks or unsupervised learning algorithms like clustering. Nevertheless, these methods typically underwent training with a limited set of micrographs. For instance, crYOLO was trained with only 840 micrographs. Consequently, they may struggle to generalize effectively to diverse protein types characterized by irregular and complex shapes, as well as heterogenous conformations. They often overlook the diversity of the proteins and are usually evaluated on one or a few simple datasets like Apoferritin and Keyhole Limpet Hemocyanin (KLH) due to lack of manually annotated particle data. Among these methods, crYOLO and Topaz are most widely used. CrYOLO utilizes the You Only Look Once (YOLO), an object detection algorithm^26^ trained on cryo-EM micrographs, and Topaz employs positive-unlabeled convolutional neural networks^20^ for particle picking. While both approaches have demonstrated significant potential in automating particle picking, their training has been based on a relatively small number of micrographs. CrYOLO often misses many true protein particles while Topaz picks too many particles including false positives and duplicates. The large number of particles picked by Topaz also causes difficulty in storing and processing the extracted particles required for the down-stream processing steps. As a result, the potential of deep learning for particle picking has not yet been fully harnessed, and the cryo-EM community still needs to mostly rely on traditional semi-automated methods like template-based picking tools like RELION and CryoSPARC to perform particle picking, which are time consuming and error-prone.

Two recent developments provide good opportunities to further improve automated particle picking. The first is the recent creation of a large, labeled protein particle dataset -CryoPPP^4^ from the Electron Microscopy Public Image Archive (EMPIAR)^27^, which enables the development and training of sophisticated deep learning methods for particle picking. The second one is the availability of large foundational AI image segmentation models such as Meta’s Segment Anything Model (SAM) ^28^ that may be used to segment objects in images. However, a direct application of SAM to cryo-EM images can segment few particles because cryo-EM images are very different from the image data used to train SAM. Moreover, a simple retraining of SAM on cryo-EM images only yielded somewhat improved but still unsatisfactory results.

To leverage the opportunities and address the challenges above, we first designed a specialized U-Net architecture^29^ with the inclusion of attention gates in each decoder block and trained it on the CryoPPP dataset to pick protein particles. After training, the attention-gated U-Net is applied to any cryo-EM micrograph to generate a segmentation map as input for SAM’s automatic mask generator^28^ for accurately localizing protein particles in the cryo-EM micrograph. This segmentation network of integrating the specialized U-Net architecture and SAM for particle picking (called CryoSegNet) performs better than the two most popular AI based pickers crYOLO and Topaz in terms of both the accuracy of particle picking and the resolution of 3D protein density maps reconstructed from picked particles. Particularly, CryoSegNet substantially increases the resolution of density maps constructed from picked particles over crYOLO and Topaz, making it a useful tool for generating more accurate protein structures from both existing and new cryo-EM image data.

## Results

### I. Combining the specialized attention-gated U-Net trained on cryo-EM images with the general foundational Segment Anything Model (SAM) for particle picking

**Fig. 1** illustrates the process of particle picking from cryo-EM micrographs using CryoSegNet. A cryo-EM micrograph is first denoised by the image processing techniques^22,30,31^. The denoised micrograph is then used as input for an attention-gated U-Net trained on a comprehensive and diverse dataset consisting of thousands of manually labeled cryo-EM micrographs of 22 diverse protein types to pick particles to generate a segmentation map, which is used as input for SAM to generate a mask map with identified particles. The particles in the mask map are further post-processed (e.g., combined or filtered) by a post-processing module to generate the final output containing the picked particles. The final output includes the protein particle coordinates in the form of .star files, which are compatible with widely used tools like RELION^11^ and CryoSPARC^12^ and can be directly used by them to generate 3D protein density maps. The design and training of the attention-gated U-Net and the details of each processing step above are described in the **Methods** section.

**Fig. 1.**
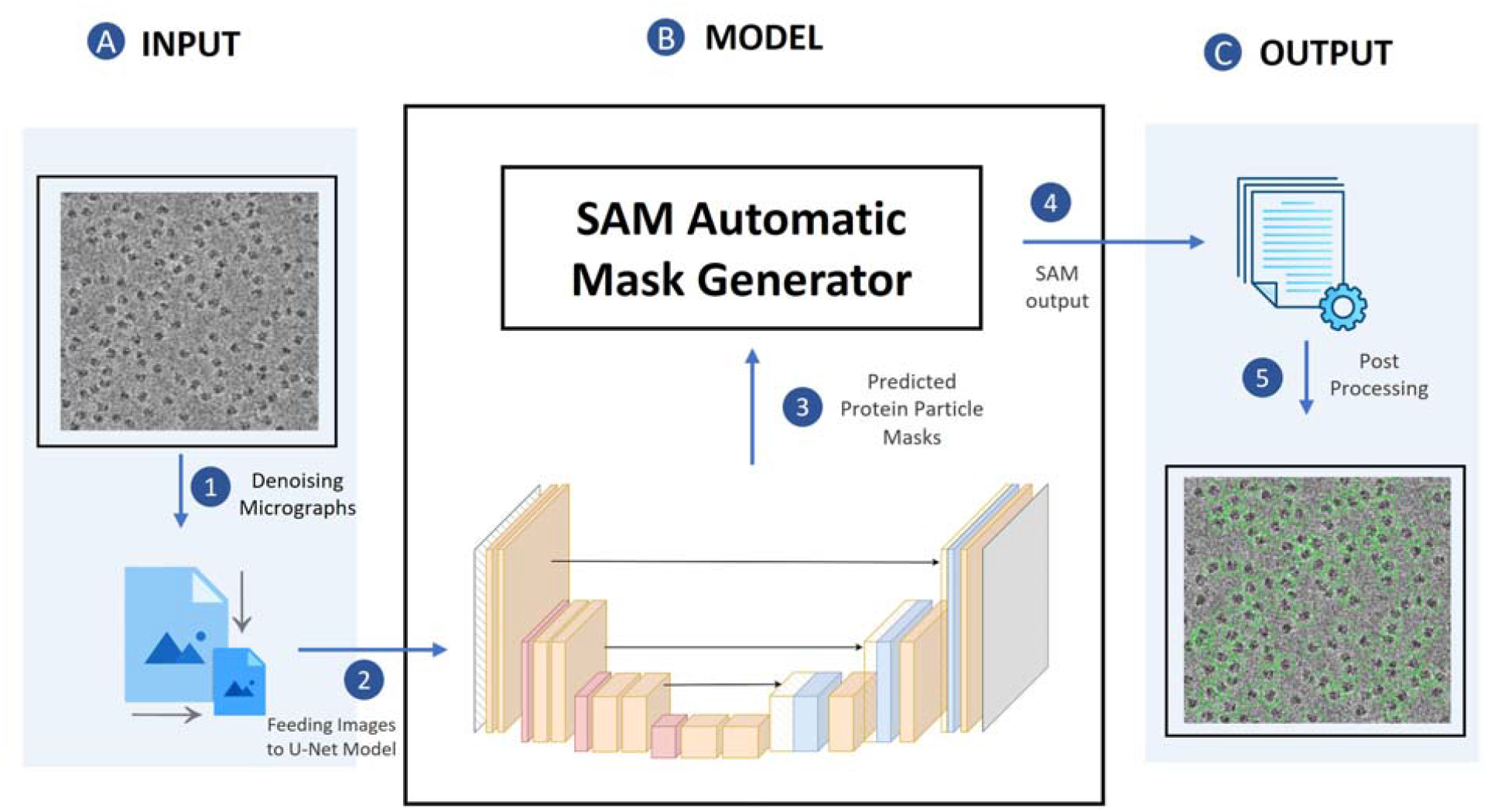
The process of particle picking with CryoSegNet. (**A)** An input micrograph is first denoised and then sent to the U-Net model. (**B)** U-Net model outputs a segmentation mask for each micrograph that is fed to SAM automatic mask generator for predicting the bounding boxes of protein particles. (**C**) The output generated by SAM is further processed based on thresholding the prediction confidence scores to filter out some false particles to generate the final output of picked particles stored in .star files.

After CryoSegNet was trained and validated on the training/validation, we blindly benchmarked it on a test dataset consisting of thousands of labeled cryo-EM micrographs of 7 different protein types from the CryoPPP^4^ dataset. The particles picked by CryoSegNet were compared with the ground truth coordinates of the expert-labeled particles.

The standard image segmentation metrics including precision, recall, F1-score 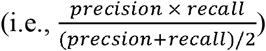, and Dice score^32^ of particle picking made by CryoSegNet were calculated to evaluate its performance. Dice score is used to evaluate the similarity between predicted segmentation masks and ground truth masks. It ranges from 0 (zero overlap) to 1 (perfect overlap). Furthermore, as an ultimate test, we constructed 3D density maps for each protein from the particles picked by CryoSegNet, crYOLO and Topaz respectively and compared the resolution of the reconstructed density maps. The detailed results are reported in the sub-sections below.

### II. The performance of particle picking on the CryoPPP test dataset in terms of image segmentation metrics

The number of cryo-EM micrographs and labeled particles for each of the seven different types of proteins in the CryoPPP test dataset is reported in **Table 1**. There are 1,879 labeled cryo-EM images and 401,263 labeled particles in total, which form the largest test dataset for evaluating particle picking methods to date. To fairly compare the three methods: CrYOLO, Topaz and CryoSegNet, we trained and tested all these methods with the same set of training, validation and test data. The CrYOLO was trained with “PhosaurusNet” architecture and Topaz with “ResNet16” architecture. The details of parameters used in training of CrYOLO and Topaz can be found in **Supplementary Note S1**. The per-protein and average precision, recall, F1-score, and Dice score of CryoSegNet, crYOLO, and Topaz on the dataset are summarized in **Table 1**. The average precision, recall, F1-score, and Dice score of CryoSegNet are 0.792, 0.747, 0.761 and 0.719 respectively, while for CrYOLO, they are 0.744, 0.768, 0.751, and 0.698. Topaz has an average precision, recall, F1-score, and Dice score of 0.704, 0.802, 0.729, and 0.683, respectively. Among the three methods, CryoSegNet has the highest F1-score, precision, and Dice score, while Topaz has the highest recall. The higher F1-score of 0.761 for CryoSegNet, in contrast to 0.729 for Topaz and 0.751 for CrYOLO, indicates that CryoSegNet is a more balanced particle picker than Topaz and CrYOLO, considering both sensitivity (recall) and specificity (precision).

**Table 1.** Evaluation results on the CryoPPP test dataset. The EMPIAR ID of the cryo-EM image set for each of the 7 test proteins is listed in Column 1. The type of each protein, number of cryo-EM images and number of labeled particles are reported in Columns 2-4. The precision, recall, F1-score, and Dice score for crYOLO, Topaz and CryoSegNet are reported in the other columns. Bold font denotes the best average score of each metric.

| EMPIAR ID | Type of Protein | Num. of Labeled Images | Num. of Labeled Particles | CrYOLO |  |  |  | Topaz |  |  |  | CryoSegNet |  |  |  |
| --- | --- | --- | --- | --- | --- | --- | --- | --- | --- | --- | --- | --- | --- | --- | --- |
|  |  |  |  | Precision | Recall | F1 Score | Dice Score | Precision | Recall | F1 Score | Dice Score | Precision | Recall | F1 Score | Dice Score |
| 10028<sup>33</sup> | Ribosome (80S) | 300 | 26,391 | 0.807 | 0.941 | 0.869 | 0.863 | 0.696 | 0.937 | 0.799 | 0.786 | 0.833 | 0.944 | 0.885 | 0.859 |
| 10081<sup>34</sup> | Transport | 300 | 39,352 | 0.822 | 0.884 | 0.852 | 0.822 | 0.732 | 0.872 | 0.796 | 0.758 | 0.835 | 0.922 | 0.876 | 0.876 |
| 10345<sup>35</sup> | Signaling | 295 | 15,894 | 0.648 | 0.665 | 0.656 | 0.452 | 0.544 | 0.805 | 0.650 | 0.507 | 0.746 | 0.920 | 0.824 | 0.743 |
| 11056<sup>36</sup> | Transport | 305 | 125,908 | 0.726 | 0.780 | 0.752 | 0.718 | 0.764 | 0.909 | 0.830 | 0.778 | 0.757 | 0.687 | 0.720 | 0.663 |
| 10532<sup>37</sup> | Viral | 300 | 87,933 | 0.756 | 0.774 | 0.765 | 0.724 | 0.732 | 0.939 | 0.823 | 0.788 | 0.796 | 0.628 | 0.702 | 0.649 |
| 10093<sup>38</sup> | Membrane | 295 | 56,394 | 0.623 | 0.744 | 0.678 | 0.641 | 0.610 | 0.216 | 0.319 | 0.279 | 0.716 | 0.515 | 0.600 | 0.537 |
| 10017<sup>39</sup> | β-galactosidase | 84 | 49,391 | 0.824 | 0.588 | 0.686 | 0.663 | 0.847 | 0.936 | 0.889 | 0.886 | 0.859 | 0.616 | 0.718 | 0.703 |
| **Average** |  |  |  | 0.744 | 0.768 | 0.751 | 0.698 | 0.704 | **0.802** | 0.729 | 0.683 | **0.792** | 0.747 | **0.761** | **0.719** |

Moreover, we compared the predictions made by the three methods for some individual micrographs to study their characteristics. **Fig. 2** illustrates the typical disparities in particle picking among crYOLO, Topaz and CryoSegNet on three individual cryo-EM micrographs of two protein types (EMPIAR ID 10345 and EMPIAR ID 11056). CrYOLO tends to pick fewer protein particles, thereby discarding many true particles. Topaz, when using with default parameters, picks an excessive number of true particles with a lot of overlaps (redundancy) as well as false particles within carbon edges and ice patches that can cause a serious difficulty for the 3D reconstruction of density maps from the picked particles. The storage requirement for processing the redundant particles from Topaz for 3D reconstruction is substantial. In contrast, CryoSegNet usually picks most true protein particles while selecting only a small number of false positives, minimizing the number of redundant/duplicated/overlapped particles and largely excluding false particles in the carbon edges and ice patches.

**Fig. 2.**
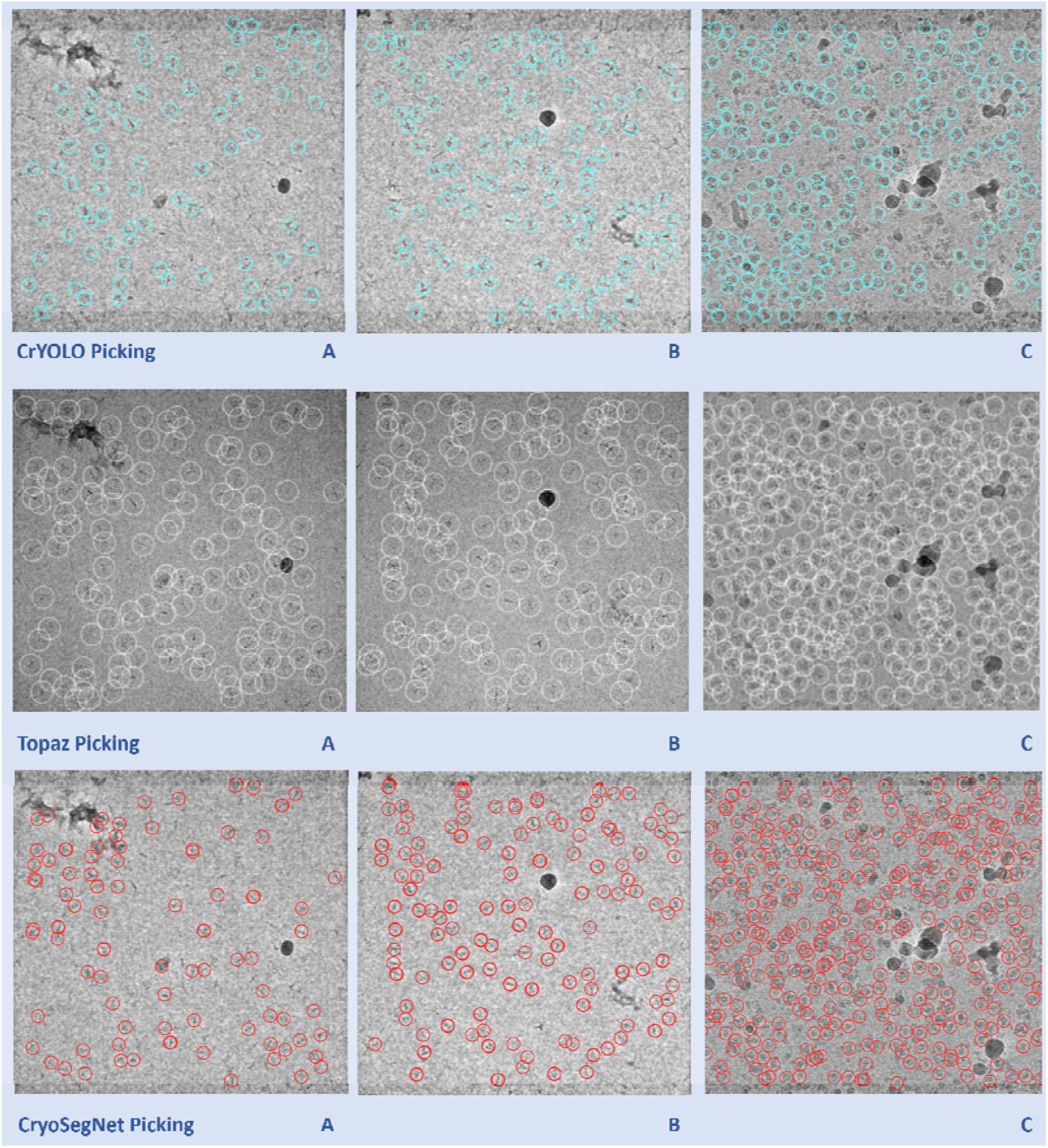
Comparison of particle picking by crYOLO, Topaz and CryoSegNet on three cryo-EM micrographs of two protein types (EMPIAR ID 10345 and EMPIAR ID 11056). **(A)** Topaz picks ice patches and more particles in the contaminated regions than CryoSegNet while crYOLO picks few particles (EMPIAR ID 10345). **(B)** Topaz picks more false positives (particularly the ones on the black ice patch) compared to CryoSegNet (EMPIAR ID 10345). **(C)** CryoSegNet picks a zero to small number of particles in undesired (carbon or ice) regions (black holes) of the micrograph (EMPIAR ID 11056), while Topaz picks some false particles in the regions.

We also compare the precision, recall, F1-score, and Dice score of the output of each of the three prediction modules of CryoSegNet: (1) the attention-gated U-Net, (2) the SAM and (3) the postprocessing module (**Supplementary Table S1**). At the end of each subsequent module, the F1-scores are computed, revealing higher values for SAM (0.768) and the postprocessing module (0.761) in comparison to U-Net (0.71). This indicates that the performance is improved by incorporating SAM into the output of U-Net. Interestingly, applying the SAM module to the output of the U-Net substantially increases the recall from 0.739 to 0.820, while decreasing the precision from 0.747 to 0.729. Adding the post-processing on top of the SAM output increases the precision from 0.729 to 0.792, while decreasing the recall from 0.820 to 0.747. At the end, the precision of the final output of CryoSegNet (e.g., the output of the post-processing module) is substantially higher than the U-Net (0.792 versus 0.747), while its recall is slightly higher than the U-Net (0.747 versus 0.739), resulting in a higher F1-score (0.761 versus 0.71). The results show that the three prediction steps of CryoSegNet complement each other, leading to the balanced performance.

To further assess the performance of these methods, we fine-tuned each of the three pre-trained methods above for each EMPIAR ID in the test dataset by using 20 labeled micrographs as training and validation data and the remaining micrographs as the test data. We compared the fined-tuned CryoSegNet with the fine-tuned CrYOLO and fine-tuned Topaz on the withheld test data of each EMPIAR ID. The overall average performance of each method is improved by the fine tuning, indicating that fine-tuning each method using a small number of human-labeled micrographs for a target protein can further enhance the accuracy of particle picking. The fine-tuned CryoSegNet still has higher F1-score, precision, and Dice score than the fine-tuned CrYOLO and Topaz. The detailed results are presented in **Supplementary Table S2**.

### III. The performance of particle picking in terms of the resolution of 3D density maps reconstructed from picked particles

The F1-score, precision and recall of particle picking can measure the accuracy of a machine learning method discriminating particles from non-particles, but they do not directly measure the quality of the density maps of proteins reconstructed from the picked particles, which are the end product concerning users most. Reconstructing 3D density maps from picked particles involves very complex algorithms of converting 2D particle images to 3D density maps, whose performance depends on many factors such as the number of true particles, the uniqueness of true particles capturing different orientations (views) of protein structure, and the severity of false particles that cannot be simply measured by a single score such as F-measure, precision and recall. Therefore, as an ultimate test, we compare CryoSegNet, Topaz, and crYOLO in terms of the resolution of 3D density maps reconstructed from picked particles on CryoPPP test dataset.

#### A. The comparison of the resolution of the density maps reconstructed from the particles picked by CryoSegNet, crYOLO and Topaz on CryoPPP test dataset

For each protein type in the test dataset, we generate star files containing particles picked by a method, which are then imported into CryoSPARC for 3D ab-initio reconstruction of density maps and homogenous refinement^12^. In the context of ab-initio reconstruction, we reconstruct a 3D density map from only a set of particles without using any initial structural model or starting structure as input. Homogeneous refinement is employed to rectify higher-order aberrations and to refine particle defocus caused by factors such as beam tilt, spherical aberration, and other optical issues. We compare the 3D resolution of the density maps reconstructed from the particles picked by crYOLO, Topaz, and CryoSegNet. Results are computed both with and without considering the best 2D templates from the Select2D job^12^ in CryoSPARC. Select2D is a process used by CryoSPARC internally to filter out low-quality/false particles provided by users before the density map reconstruction.

The experiments were conducted across three trials with random seed initialization, and the best resolution was considered for comparison. The summary results of the three methods on the micrographs in CryoPPP test dataset are presented in **Table 2**, while the detailed trial results can be found in **Supplementary Table S3**. The resolution of both CryoSegNet and Topaz is higher than crYOLO on 6 out of 7 protein types. CryoSegNet has a higher resolution than Topaz on 5 out of 7 protein types and a lower resolution than Topaz on two protein types. The average resolution of CryoSegNet with Select 2D is 4.94 Å, better than 5.16 Å of Topaz and 5.29 Å of crYOLO. Also, on all 7 protein types, Topaz picked most particles (67,906 on average), CryoSegNet second most (46,893 on average), and crYOLO least (42,475 on average), indicating that the quality of density maps does not fully depend on the number of picked particles. This result can be largely explained by the observation that crYOLO picks fewer particles, Topaz identifies many particles with some redundancy/overlap, and CryoSegNet picks most true particles with little redundancy.

**Table 2.** Comparison of CryoSegNet with crYOLO and Topaz in terms of the resolution of 3D density maps on CryoPPP test dataset. Bold font denotes the highest resolution.

| EMPIAR ID | Without Select 2D |  |  |  |  |  | With Select 2D |  |  |  |  |  |
| --- | --- | --- | --- | --- | --- | --- | --- | --- | --- | --- | --- | --- |
|  | Number of Picked Particles |  |  | Best Resolution (Å) |  |  | Number of Particles |  |  | Best Resolution (Å) |  |  |
|  | CrYOLO | Topaz | CryoSegNet | CrYOLO | Topaz | CryoSegNet | CrYOLO | Topaz | CryoSegNet | CrYOLO | Topaz | CryoSegNet |
| 10028 | 32,687 | 52,588 | 47,764 | 4.13 | 3.98 | **2.72** | 31,699 | 35,514 | 45,218 | 4.11 | 3.93 | **2.72** |
| 10081 | 44,440 | 58,217 | 60,158 | 5.65 | 6.13 | **4.58** | 36,821 | 37,808 | 44,819 | 4.97 | 5.08 | **4.16** |
| 10345 | 15,821 | 29,208 | 25,919 | 3.98 | 3.73 | **3.48** | 11,369 | 21,343 | 15,209 | 3.83 | 3.64 | **2.84** |
| 11056 | 60,648 | 98,680 | 71,342 | 8.98 | 8.11 | **7.83** | 43,599 | 66,651 | 53,073 | 8.32 | 8.03 | **7.13** |
| 10532 | 46,162 | 73,196 | 67,219 | 4.23 | 4.54 | **4.09** | 29,434 | 38,372 | 30,155 | 4.08 | 4.23 | **3.89** |
| 10093 | 43,305 | 110,577 | 43,886 | 7.27 | **6.35** | 7.27 | 33,183 | 61,698 | 27,745 | 6.87 | **6.12** | 6.99 |
| 10017 | 54,263 | 52,875 | 11,961 | **4.99** | 5.13 | 6.90 | 47,704 | 45,511 | 10,026 | **4.84** | 5.08 | 6.86 |
| **Average** | 42,475 | 67,906 | 46,893 | 5.60 | 5.42 | **5.27** | 33,401 | 43,842 | 32,321 | 5.29 | 5.16 | **4.94** |

Moreover, applying Select 2D to the density map reconstruction improves the resolution of all these methods. It is worth noting that, even though the results in **Table 2** were obtained from particles picked from at most 305 micrographs for each protein type in CryoPPP test dataset, the resolution of CryoSegNet for some protein types is high. For instance, on two protein types (EMPIAR ID 10028 and 10345), the resolution of CryoSegNet, after removing some false positives by Select 2D, is below 3 Å.

#### B. The comparison of resolution of 3D density maps reconstructed from all cryo-EM micrographs of five protein types in EMPIAR

In addition to evaluating the on the test dataset from CryoPPP that has only approximately 300 micrographs for each protein type (see **Table *1***), we extended the assessment of the methods to the complete set of micrographs available on the EMPIAR website for five different protein types in CryoPPP test dataset (**Table *3***) to gauge the resolution that they can achieve in a real-world setting. CryoSegNet and Topaz substantially outperform crYOLO on each protein type and on average. Moreover, CryoSegNet performs better than Topaz for all the protein types except EMPIAR ID 10093. The average resolution of CryoSegNet with Select 2D is 3.32 Å, about 7% better than 3.57 Å of Topaz and 14% better 3.85 Å of crYOLO. Remarkably, for EMPIAR ID 10345, the resolution of the density map reconstructed from CryoSegNet is 2.67 Å, which is much higher than CrYOLO and Topaz. Moreover, the average resolution across all test sets resulting from CryoSegNet picked particles (3.32 Å) is 3% better than the average 3.33Å of the density maps built by their original authors possibly with some manual particle picking, and CryoSegNet has a better resolution than the original ones for three out of five proteins, indicating that it can be applied to the existing cryo-EM micrographs in EMPIAR to generate better density maps.

**Table 3.** Comparison of 3D resolution of on the full set of micrographs of five protein types. The last column lists the resolution of the density maps built by their original authors as a reference.

| EMPIAR ID | Without Select 2D |  |  |  |  |  | With Select 2D |  |  |  |  |  | Original EMPIAR Resolution (Å) |
| --- | --- | --- | --- | --- | --- | --- | --- | --- | --- | --- | --- | --- | --- |
|  | Number of Particles |  |  | Best Resolution (Å) |  |  | Number of Particles |  |  | Best Resolution (Å) |  |  |  |
|  | CrYOLO | Topaz | CryoSegNet | CrYOLO | Topaz | CryoSegNet | CrYOLO | Topaz | CryoSegNet | CrYOLO | Topaz | CryoSegNet |  |
| 10028 | 65,376 | 104,652 | 93,881 | 3.97 | **2.72** | **2.72** | 63,562 | 96,352 | 92,532 | 3.94 | **2.72** | **2.72** | 3.20 |
| 10345 | 50,506 | 102,977 | 120,357 | 3.56 | 3.50 | **2.74** | 40,047 | 87,472 | 73,377 | 3.54 | 3.45 | **2.67** | 3.51 |
| 10081 | 148,488 | 171,396 | 202,988 | 4.26 | 4.34 | **3.95** | 123,963 | 130,941 | 153,333 | 4.15 | 4.06 | **3.45** | 3.50 |
| 10532 | 232,220 | 362,115 | 181,259 | 3.25 | 3.52 | **3.42** | 161,497 | 206,460 | 90,477 | 3.22 | 3.22 | **3.20** | 2.90 |
| 10093 | 264,447 | 801,208 | 267,983 | **4.54** | 4.55 | 4.70 | 192,337 | 437,235 | 169,330 | 4.41 | **4.40** | 4.54 | 3.55 |
| Average | 152,207 | 308,470 | 173,294 | 3.92 | 3.73 | **3.51** | 116,281 | 191,692 | 115,810 | 3.85 | 3.57 | **3.32** | 3.33 |

Comparing the results on all the micrographs of the five protein types (**Table 3**) and the results on a smaller number of micrographs of the same five protein types (**Table 2**), the average performance of all three methods on the five protein types is improved, indicating that using more micrographs generally improve the quality of reconstructed density maps as expected. Moreover, applying Select 2D to the density map reconstruction improves the resolution of all the three methods on this dataset, even though Select 2D filters out a substantial number of particles including some true ones picked by each method, indicating that other factors such as the quality and representativeness of picked particles are important. This explains why a single particle picking metric such as recall (sensitivity) does not fully correlate with the resolution of reconstructed density maps. The detailed results of the three methods in all the trials can be found in **Supplementary Table S4**.

The superiority of CryoSegNet is not only evident in terms of resolution but also in the quality of viewing direction and the representation of various orientations of picked particles. **Fig. 3** showcases the best 2D classes for the five protein types obtained from CryoSegNet, which clearly shows that CryoSegNet picked particles representing many different orientations/views of proteins, which is an important factor of obtaining high-resolution reconstruction of 3D density maps. Further,

**Fig. 3.**
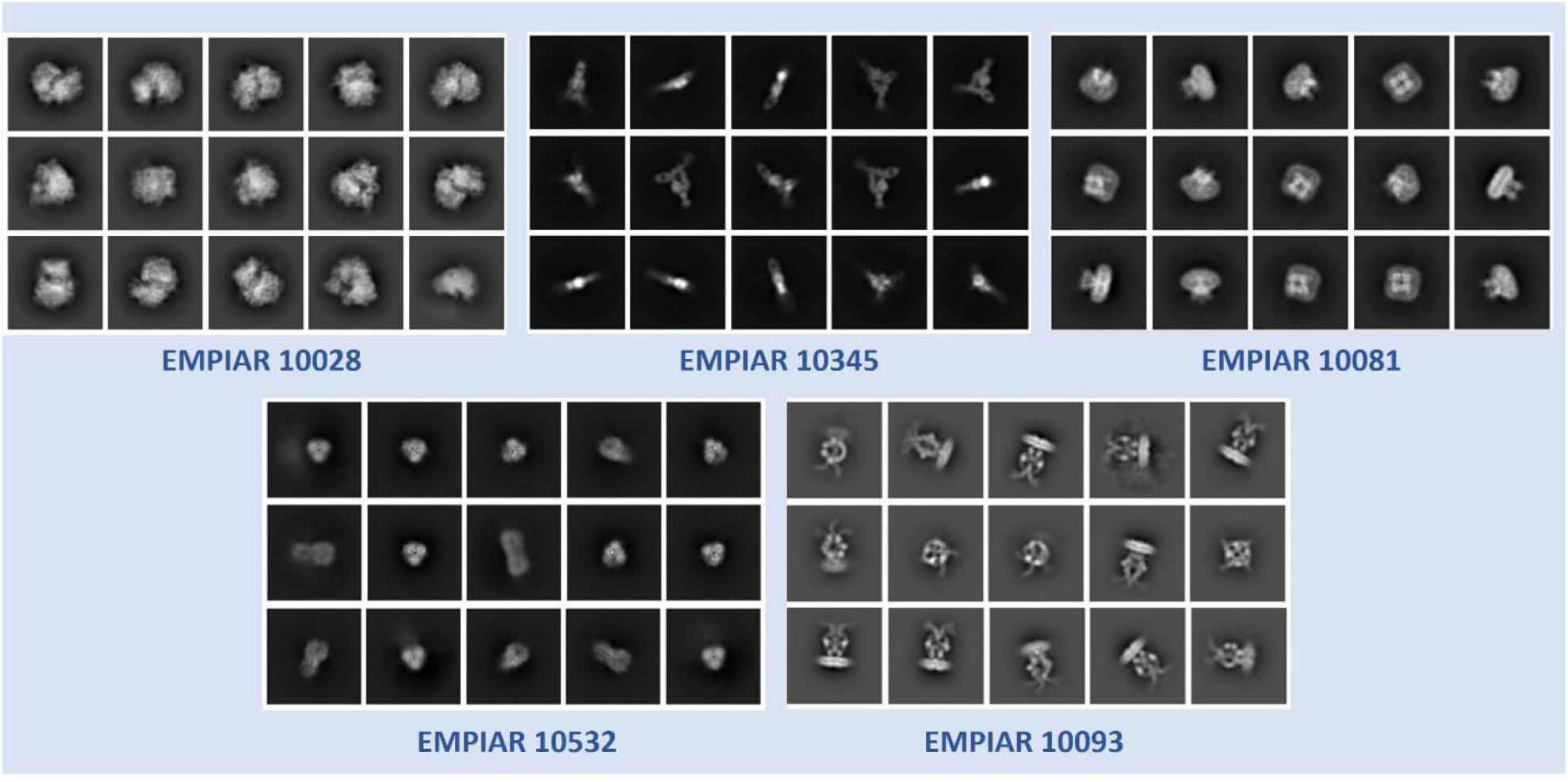
2D classes from particles picked by CryoSegNet for EMPIAR 10081, EMPIAR 10345, EMPIAR 10532, EMPIAR 10028 and EMPIAR 10093. These classes show particles with multiple orientations that have been picked by CryoSegNet.

**Fig. *4*** illustrates the comparison of viewing direction, resolution, and 3D density map of the particles picked by crYOLO, Topaz and CryoSegNet, visually showing that CryoSegNet performs better than crYOLO for all the protein types and better than Topaz for most protein types.

**Fig. 4.**
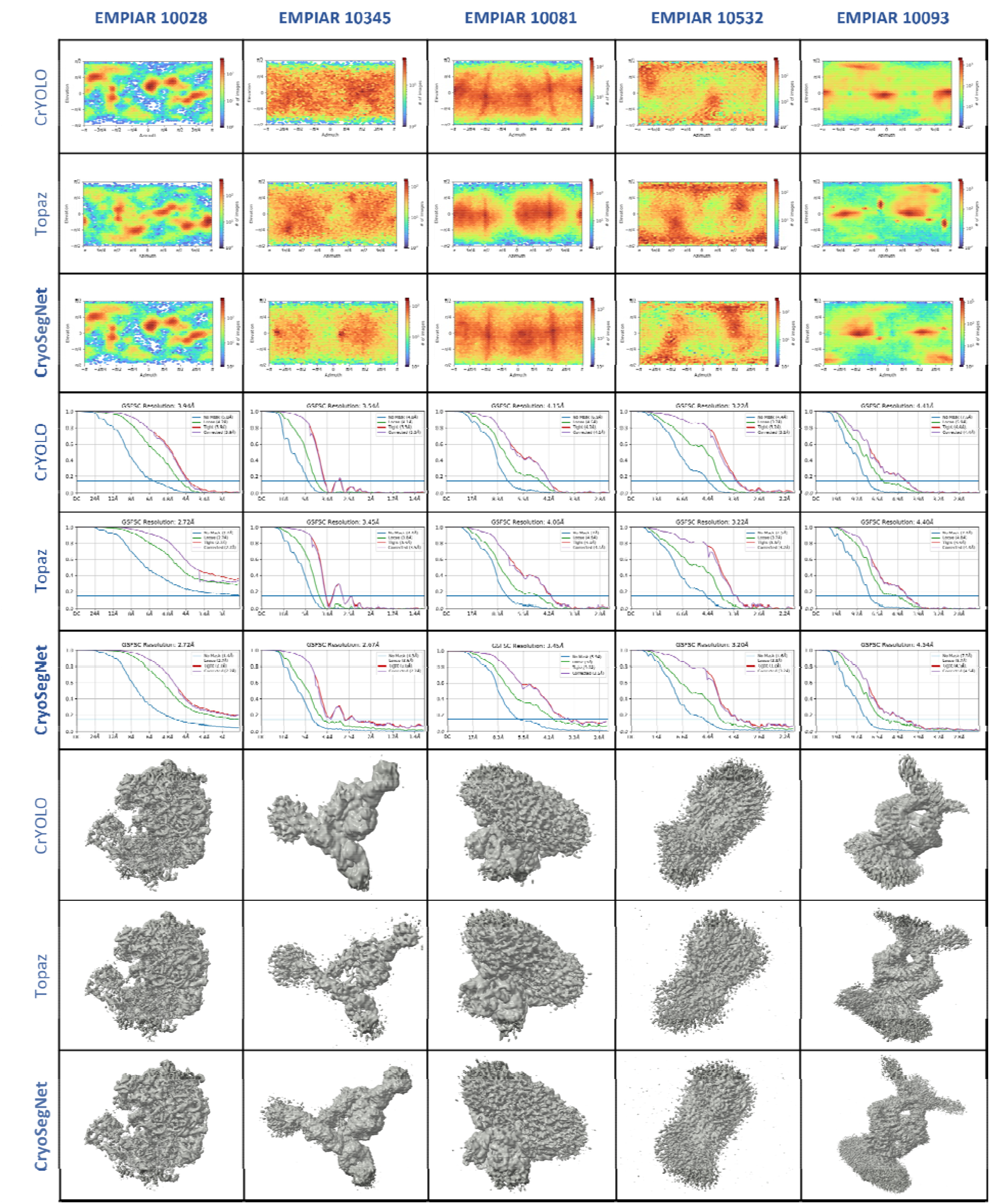
Comparison results for viewing direction, resolution, and 3D density map of particles picked by crYOLO, Topaz and CryoSegNet. The top 3 rows illustrate the viewing direction comparison, the middle 3 rows show the resolution comparison, and the bottom 3 rows illustrate the 3D density map comparison. From the viewing direction plots, it is observed that crYOLO picks very few particles and misses many true protein particles and CryoSegNet picks particles with multiple orientations/views. 3D density maps for CryoSegNet have much better resolution and low noise compared to crYOLO in all the cases and better resolution than Topaz for most of the protein types.

#### C. How does the resolution of density maps change with respect to the number of micrographs?

We further analyzed the impact of the number of micrographs on the resolution of the reconstructed 3D density maps for the five protein types by comparing the performance of CryoSegNet on a few hundred micrographs in CryoPPP test dataset and the full set of micrographs in EMPIAR (**Table 4**). The results show that augmenting the number of micrographs generally results in an increased number of protein particles at different viewing directions on four of five protein types, thereby contributing to improved resolution. However, if the number of particles with different conformations remains unchanged, increasing the number of micrographs does not significantly impact the final 3D resolution. For example, EMPIAR ID 10028 (ribosome), the resolution of using 300 micrographs is 2.72 Å, which is the same as that of using 600 micrographs.

**Table 4.** Comparative analysis of 3D resolution of CryoSegNet between the complete EMPIAR micrograph set and the smaller CryoPPP test dataset.

| EMPIAR ID | CryoPPP Dataset |  |  | EMPIAR Dataset |  |  |
| --- | --- | --- | --- | --- | --- | --- |
|  | Number of Micrographs | Number of Particles | Best Resolution (Å) | Number of Micrographs | Number of Particles | Best Resolution (Å) |
| 10028 | 300 | 47,764 | **2.72** | 600 | 92,532 | **2.72** |
| 10345 | 295 | 25,919 | 2.84 | 1,644 | 73,377 | **2.67** |
| 10081 | 300 | 60,158 | 4.16 | 997 | 153,333 | **3.45** |
| 10532 | 300 | 67,219 | 3.89 | 1,556 | 90,477 | **3.20** |
| 10093 | 295 | 43,886 | 6.99 | 1,873 | 169,330 | **4.54** |

The results show that the pretrained CryoSegNet has the ability to pick protein particles in large new datasets with great accuracy leading to high resolution density maps. Moreover, for some protein types, fine-tuning the pretrained model can lead to even better results. We fine-tuned CrYOLO, Topaz and CryoSegNet for each EMPIAR ID in the EMPIAR test dataset and compared the results. The details of the fine-tuning can be found in **Supplementary Note S2** and the results are shown in **Supplementary Table S5** and **Supplementary Table S6**.

### IV. Comparison of CryoSegNet and CASSPER

To further assess the performance of CryoSegNet, we compared it with another segmentation method -CASSPER and compared their results on the micrographs of 3 different proteins, EMPIAR ID 10017, 10081 and 10089, which were used to train, validate, and test the CASSPER model. EMPIAR ID 10017, 10081 are in the CryoPPP test dataset, while EMPIAR ID 10089 does not exist in the CryoPPP dataset at all. We fine-tuned both the pretrained CASSPER model and pretrained CryoSegNet model with 20 micrographs from each of these three datasets as training and validation data and then tested them on the remaining micrographs (test datasets). We then compared the 3D resolution of density maps reconstructed from the particles in the test datasets picked by the two fine-tuned models. The details of the resolution of the reconstructed density maps and the number of particles picked are presented in **Table 5**. CryoSegNet performs better than CASSPER on each of the three datasets. CryoSegNet has an average resolution of 4.04 Å, better than 4.53 Å of over CASSPER.

**Table 5.** Comparison of CryoSegNet with CASSPER in terms of the resolution of 3D density maps. Bold font denotes the highest resolution.

| EMPIAR ID | Number of Micrographs | CASSPER |  |  |  | CryoSegNet |  |  |  |
| --- | --- | --- | --- | --- | --- | --- | --- | --- | --- |
|  |  | Number of Particles | Best Resolution Å (Without Select 2D) | Number of Particles | Best Resolution Å (With Select 2D) | Number of Particles | Best Resolution Å (Without Select 2D) | Number of Particles | Best Resolution Å (With Select 2D) |
| 10017 | 84 | 44,213 | 5.38 | 38,460 | 5.32 | 38,349 | 5.27 | 31,941 | **5.2** |
| 10081 | 997 | 133,366 | 4.37 | 115,297 | 4.18 | 202,988 | 3.95 | 153,333 | **3.56** |
| 10089<sup>40</sup> | 97 | 14,565 | 4.37 | 10,335 | 4.09 | 13,533 | 4.36 | 11,847 | **3.36** |
| Average | 393 | 64,048 | 4.71 | 54,697 | 4.53 | 84,957 | 4.53 | 65,707 | **4.04** |

## Discussion

In this study, we have introduced CryoSegNet, a novel approach for protein particle picking from cryo-EM micrographs. The results show that CryoSegNet consistently outperforms the existing particle pickers in terms of the accuracy (i.e., F1-score) of particle picking and the resolution of reconstructed 3D density maps. Particularly, it substantially outperforms the state-of-the-art deep learning particle picking method Topaz in terms of the resolution of density maps reconstructed from picked particles. These advances mostly come from two sources. The first is to train CryoSegNet on the large, comprehensive and diverse dataset for protein particle picking – CryoPPP. The second is to combine the power of multiple useful techniques, including the image processing techniques of denoising input cryo-EM micrographs, the special attention-gated U-Net for particle picking, the foundational AI model SAM, and the post-processing of the output from SAM. Combining these techniques together in CryoSegNet works better than using only one or some of them. For instance, the U-Net reduces the noise from the original cryo-EM micrographs while preserving the fine details so that the segmentation maps from the U-Net model are better understood by the SAM model for improving particle picking. The postprocessing module eliminates some of the low-quality particles and false positives generated by SAM, leading to the improved resolution of the reconstructed density maps. A detailed ablation study of the performance of pretrained SAM, fine-tuned SAM, and U-Net + SAM in particle picking is presented in **Supplementary Note S3**, demonstrating that combining the U-Net with SAM outperforms the pretrained SAM and fine-tuned SAM.

As cryo-EM particle picking is still a young field, the metrics of evaluating its performance have not been well established. In this work, we use the standard image classification metrics including precision, recall, F1-score and Dice score as well as the specialized evaluation metrics such as the resolution of density maps reconstructed from picked particles that users care about most. Each classification metric is an indicator of the performance of the particle picking but none of them is 100% correlated with the resolution of density maps. The correlation between each of the classification metrics (F1-score, precision, Dice score, and recall) and the resolution value (quality) of the density maps reconstructed from CryoSegNet with Select 2D is -0.88, -0.94, -0.91, and -0.78. The correlation is computed from the classification metric values for five protein types from **Table 1** and the resolution values of the density maps in **Table 3**. The correlation shows that the F1-score, precision and Dice score are the rather informative classification metric for predicting the quality of reconstructed density maps, which have a much stronger correlation with the resolution of the reconstructed density maps than recall. The recall is the least informative probably because when there are enough picked particles, the quality or the representativeness of the particles may be more important and low-quality or false particles may severely reduce the quality of the reconstructed density maps. Moreover, none of the standard classification metric can perfectly predict the resolution of the reconstructed density maps because the density reconstruction process is very complicated, and its outcome depends on many factors such as the quality and diversity of true particles picked that the standard classification metrics cannot measure. Therefore, the resolution of the reconstructed cryo-EM density maps is the most important metric of assessing the effectiveness of a particle picking method.

In comparison to the conventional approaches, such as manual picking and template-based methods, CryoSegNet offers a more reliable and automated solution, eliminating the need for labor-intensive manual particle selection. This presents a significant improvement in the field by minimizing human bias and increasing objectivity in particle picking. Moreover, the average resolution of the density maps reconstructed from the particles picked by CryoSegNet is higher than that of the density maps built by the original authors probably with some human intervention, indicating that CryoSegNet has the potential to substitute the time-consuming manual or template-based picking. Compared to two automated machine learning methods crYOLO and Topaz, CryoSegNet substantially improves the resolution of reconstructed density maps, indicating it can be applied to generate more accurate protein structures from the existing cryo-EM data processed by Topaz and crYOLO before or new cryo-EM data. Moreover, in terms of F1-score, precision and Dice score of particle picking – the three metrics that have the strongest correlation with the resolution of reconstructed density maps, CryoSegNet also outperforms crYOLO and Topaz.

There are still some challenges faced by AI-based particle picking methods including CrYOLO, Topaz and CryoSegNet on some datasets like EMPIAR ID 10532 and EMPIAR ID 10093 that have few samples representing rare protein view orientations, some of which could be missed by the automated AI methods. In the two cases, they performed worse than the blob-based picking^11,13,39^ in RELION used by the original authors (Table 3). One reason is that the blob-based picking was used by the authors to capture rare but diverse protein-like objects, even though it might also pick undesired false particles that required subsequent steps of false positive removal. We tested if providing a small number of labeled micrographs to fine-tune CrYOLO, Topaz, and CryoSegNet can help them identify more particles with rare view orientations. Fine-tuning the pretrained model with 20 annotated micrographs resulted in an improved resolution of density maps for CrYOLO and CryoSegNet on EMPIAR-10093 (specifically 0.3Å improvement for CrYOLO and 0.56Å improvement for CryoSegNet), but not for Topaz. On EMPIAR-10532, the fine-tuning did not improve the performance of any of the three methods, emphasizing the need for additional diverse particle samples, particularly those representing rare views, to enhance the AI-based particle picking. This analysis underscores the intricate relationship between dataset complexity, sample diversity, and algorithm efficacy in the cryo-EM particle picking. Another limitation is the requirement of high computing resources for training CryoSegNet on large cryo-EM datasets. We will explore better optimization techniques to address this issue in the future.

## Methods

### 1. Dataset

We employed an extensive and diverse dataset (CryoPPP) to train, validate and test CryoSegNet. Specifically, we utilized the micrographs of 22 EMPIAR IDs (protein types) from the CryoPPP for training and validation. We allocated 80% of the micrographs from each of the 22 protein types for training and the remaining 20% for validation. For the independent test, we selected a separate set of 7 different EMPIAR IDs from the CryoPPP dataset. The selection of EMPIAR IDs for training and testing was carefully conducted, taking into consideration various factors such as protein type, shape, size, and total structural weight. We included proteins from different categories, including transport proteins, membrane proteins, signaling proteins, viral proteins, ribosomes, aldolase, and others, each characterized by distinct shapes such as rod and circular, as well as a wide range of structural weights spanning from 77 kDa to 2198 kDa. We used a large number of cryo-EM micrographs unlike most existing machine learning methods in the field trained on very limited and simplified datasets with a small number of protein types and shapes. Our training dataset consisted of 4,948 micrographs, while our validation set was comprised of 1,244 micrographs. The details of the training dataset and validation dataset are presented in **Table 6**, while those of the independent test dataset are described in **Table 7**.

**Table 6.**
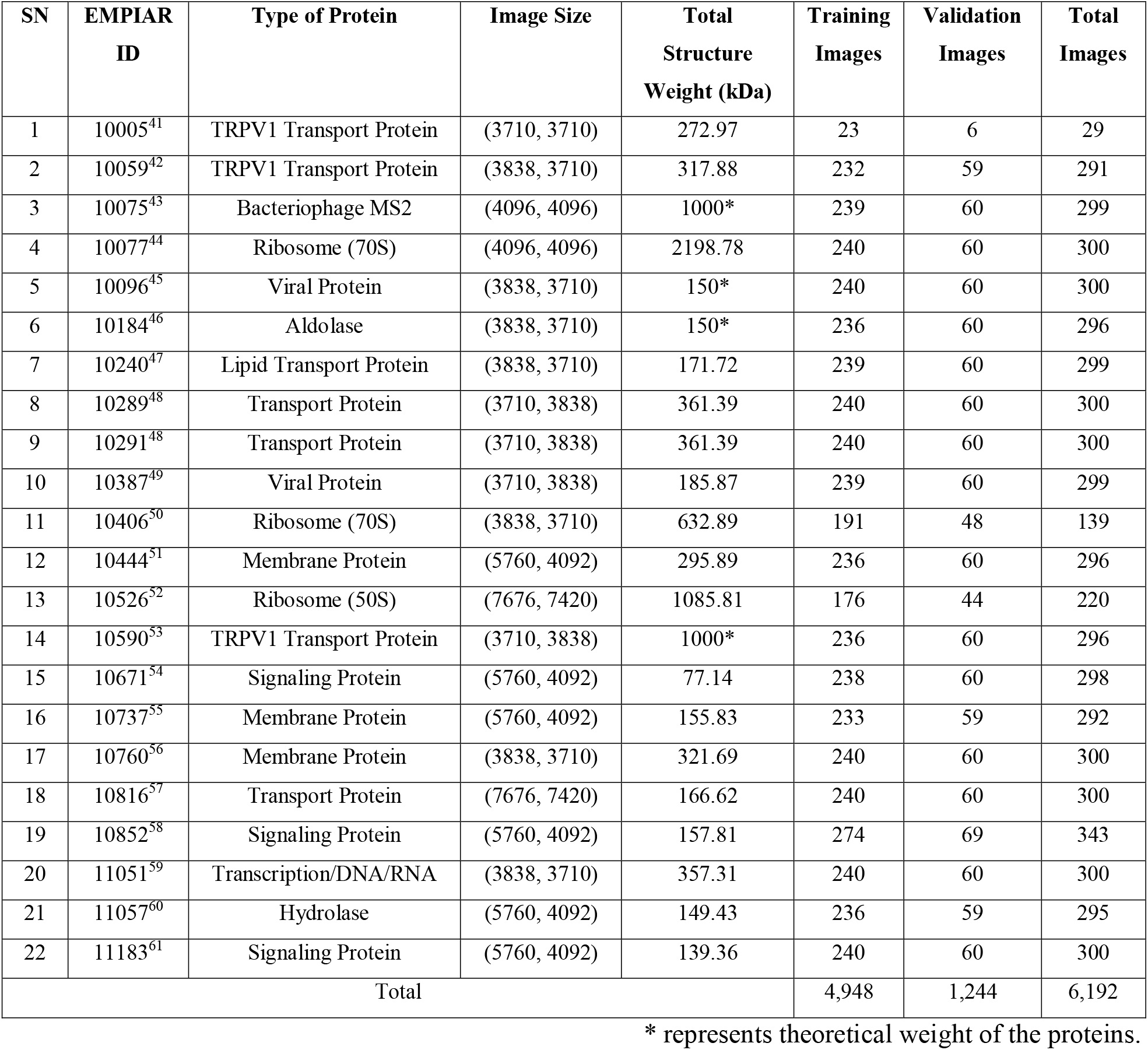
An overview of the dataset used for training and validation of CryoSegNet.

**Table 7.** An overview of the independent dataset for testing CryoSegNet.

| SN | EMPIAR ID | Type of Protein | Image Size | Total Structure Weight (kDa) | Number of Images |
| --- | --- | --- | --- | --- | --- |
| 1 | 10028 | Ribosome (80S) | (4096, 4096) | 2135.89 | 300 |
| 2 | 10081 | Transport Protein | (3710, 3838) | 298.57 | 300 |
| 3 | 10345 | Signaling Protein | (3838, 3710) | 244.68 | 295 |
| 4 | 11056 | Transport Protein | (5760, 4092) | 88.94 | 305 |
| 5 | 10532 | Viral Protein | (4096, 4096) | 191.76 | 300 |
| 6 | 10093 | Membrane Protein | (3838, 3710) | 779.4 | 295 |
| 7 | 10017 | β-galactosidase | (4096, 4096) | 450* | 84 |
| Total |  |  |  |  | 1,879 |

\* represents theoretical weight of the proteins.

### 2. Prediction Methods

#### 2.1 Attention-Gated U-Net

The advent of deep learning architectures like U-Net has greatly simplified segmentation tasks in biomedical images like localizing mitochondria cells and brain tumors. In this work, we designed a special U-Net architecture (**Fig. 5A**) for cryo-EM protein particle picking by making it deeper and introducing an attention mechanism into it, considering the large size of the cryo-EM micrographs and the nature of protein particles in the micrographs. Cryo-EM micrographs often contain objects that are not actual single protein particles, such as ice patches, protein aggregates, and false particles along the carbon edges. These false positives can negatively degrade the resolution of the final 3D structures reconstructed from the particles. Therefore, it is important to prioritize the picking of true protein particles for an accurate segmentation. Thus, we added attention gates in the expanding path of the U-Net architecture to put a significant emphasis on true protein particles. Our model consists of 5 encoder blocks in the contracting path, a bottleneck layer and 5 decoder blocks in the expanding path, each equipped with attention gates. This architecture modification can effectively handle the complexity of cryo-EM micrographs and achieve the precise segmentation of protein particles.

**Fig. 5.**
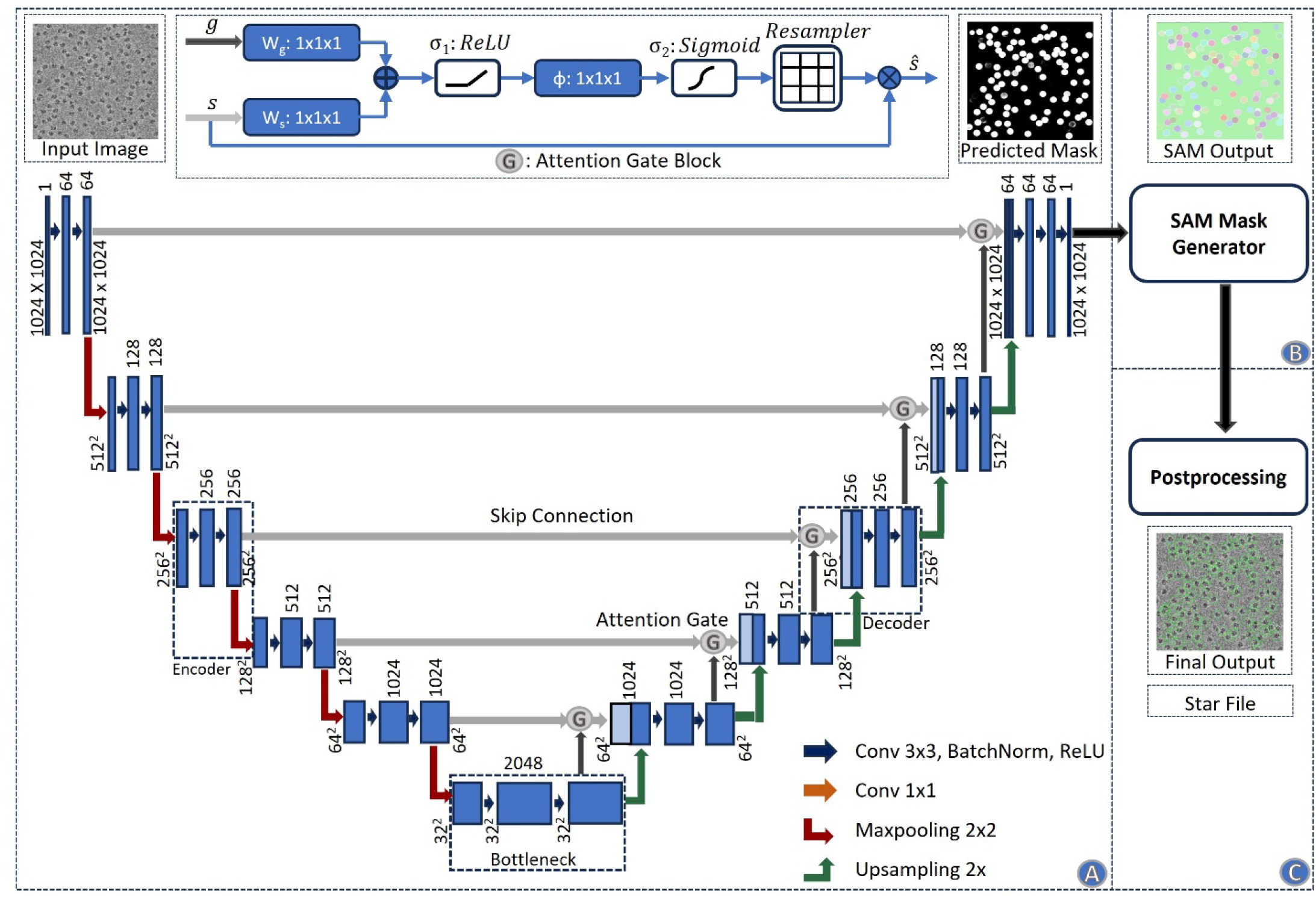
Architecture of the CryoSegNet model. **(A)** The attention-gated U-Net to predict segmentation mask for a micrograph. The numbers in the top of the rectangular slices indicate the number of channels and in the bottom indicate the size of the output. The U-Net has five encoders, one bottleneck component, and five decoders. The skip connection from each encoder to its corresponding decoder goes through an attention gated block. Each attention block for a decoder also takes an input from its previous decoder or the bottleneck component. The details of the attention block are illustrated at the middle top. **(B)** The SAM mask generator takes input from the output of the U-Net model and outputs bounding box coordinates and intersection over union score for each predicted protein particle in the micrograph. **(C)** The postprocessing module outputs the star file containing picked particles and processed output micrographs based on the thresholding criterion for each protein type.

The U-Net takes as input a cryo-EM micrograph of size 1024x1024 and outputs a segmentation mask of size 1024x1024. A loss function which combines both binary cross entropy loss and dice loss is used to measure prediction error in training. The former allows for measuring individual pixel error independently while the latter assesses the degree of dissimilarity between the predicted segmentation mask and the ground truth segmentation masks. By minimizing these two, the network is trained to achieve more accurate segmentation of protein particles. The output of the U-Net is used as input for SAM’s automatic mask generator for further segmentation.

#### 2.2 SAM automatic mask generator

Meta’s Segment Anything Model (SAM) has achieved great success in segmenting objects in many images. However, directly applying the pretrained SAM to cryo-EM micrographs can only pick very few particles because cryo-EM images are very different from the images used to train SAM. Fine tuning (retraining) the SAM’s mask decoder on cryo-EM micrographs for thousands of epochs improved results over the original SAM but still could not achieved satisfactory results and performed worse than the state-of-the-art deep learning particle pickers such as Topaz. After many trials, we finally devised a hybrid approach that combines the U-Net model with SAM’s automatic mask generator, which is proved to be highly effective for particle picking.

In the hybrid approach, the output of the attention-gated U-Net is fed to the SAM’s automatic mask generator module. This module was tailored for automatic mask generation for input images and was trained on the SA-1B dataset. Firstly, it generates the masks from a grid of points, incorporating various scales of the original and zoomed images. Then, cropping is performed using a regular grid of points, and any masks intersecting crop boundaries are discarded. Redundant masks are then eliminated through non-maximum suppression with an intersection over union (IoU) threshold of 0.7, retaining only masks with confidence scores exceeding 88.0. Subsequent processing steps refine the masks by removing small artifacts and filling minor gaps, which are particularly important considering the high noise and low contrast characteristics of cryo-EM micrographs. These refined masks as well as the IoU scores and bounding box coordinates for each picked protein particle within the micrographs are then passed through our postprocessing modules below designed to filter out some false positives and improve the precision of particle picking.

#### 2.3 Postprocessing

The output generated by SAM’s automatic mask generator undergoes the additional postprocessing to generate .star files, which contain coordinate information for protein particles. **Algorithm 1** outlines the complete steps of the postprocessing.

**Algorithm 1**. Postprocessing of the output of SAM

**Require:** a segmentation mask from SAM’s automatic mask generator as input

1. Consider only the particles with a predicted IoU greater than 0.94.

2. Extract the bounding-box information ‘bbox’ for each picked particle in the segmentation mask, where the 1^st^ and 2^nd^ values are the x and y coordinates, and the 3^rd^ and 4^th^ values are the width and height, respectively.

3. Calculate the mode of the widths (m_w) and mode of the heights (m_h) for the particles from step 2 for each segmentation mask.

4. Determine the new diameter (d) of the picked particles from each segmentation mask. Rescale the m_w and m_h values from step 3 according to the size of original micrograph. Calculate d using the formula:

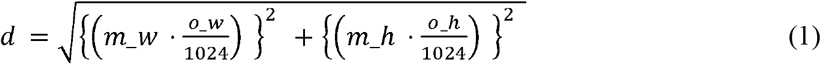

where, o_w and o_h are the width and height of the original micrograph.

5. Set a threshold value (th) equal to 10% of the diameter:

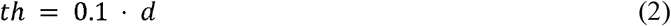

6. Select particles with width and height that satisfy the following criteria:

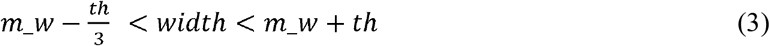

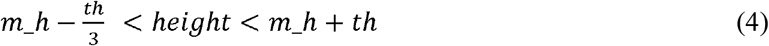

7. Calculate the scaled x and y-coordinates of the center of the protein particles for each segmentation mask of micrograph:

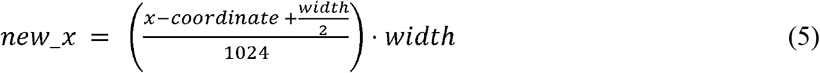

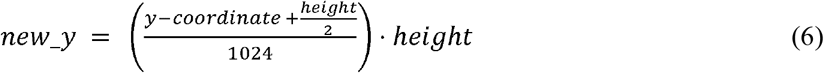

8. Output the values new_x, new_y and d of each particle from micrographs to a .star file.

### 3. Data preprocessing

#### 3.1 Denoising of micrographs

The cryo-EM micrographs have low contrast and low SNR, necessitating the use of image denoising techniques before using them as input for the U-Net. **Fig. 6** illustrates the denoising techniques used for preprocessing cryo-EM micrographs. The image preprocessing pipeline begins with reading the images in the .mrc format and applying a Gaussian filter. Subsequently, the images are standard normalized and converted to grayscale, with pixel values ranging from 0 to 255. To effectively reduce noise while preserving image details, the Fast Non-Local Means (FastNLMeans) denoising technique^22^ is applied, followed by noise mitigation through Weiner filtering^22^. To enhance the contrast of cryo-EM micrographs and improve the visibility of protein particles, the contrast limited adaptive histogram equalization (CLAHE) technique is then incorporated. CLAHE technique is widely used to enhance images with regions of non-uniform illumination and low contrast. Finally, the CLAHE equalized image is used as a guided image to the Weiner filtered image to perform guided filtering, allowing selective smoothing and enhancement of the cryo-EM micrographs while preserving edges and fine details.

**Fig. 6.**
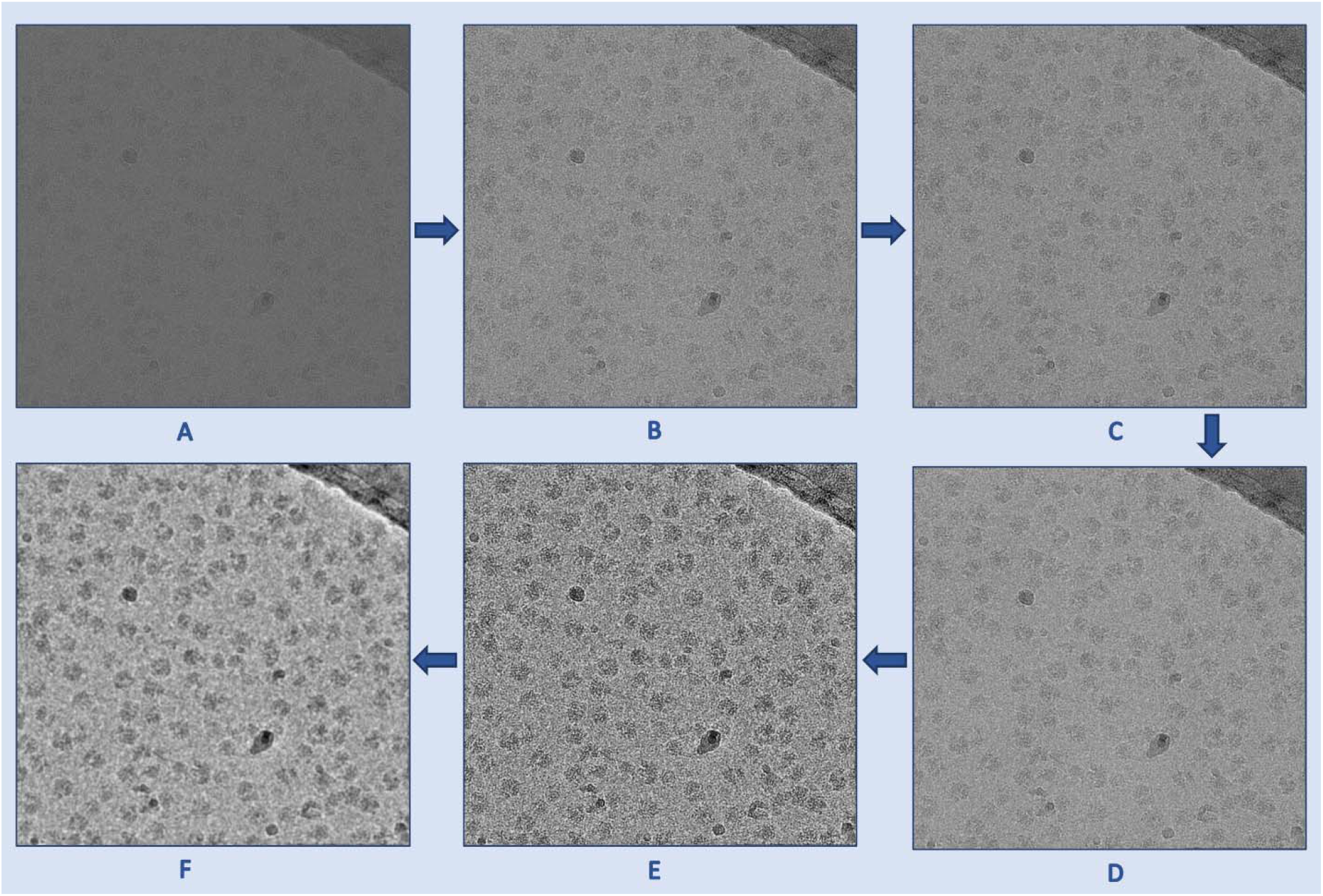
The denoising process used to preprocess cryo-EM micrographs. **(A)** An original low contrast and low SNR cryo-EM micrograph (EMPIAR ID 10406). **(B)** A standard normalized cryo-EM image. **(C)** A denoised image using FastNLMeans technique. **(D)** Weiner filter applied to the (C) for further denoising. **(E)** Contrast enhancement using CLAHE technique. **(F)** Guided filtered image with (E) as a guided image to the Weiner filtered image (D). As shown in these images, the preprocessing techniques gradually improve the contrast and SNR of the micrograph.

#### 3.2 Standardization of inputs and labels

The CryoPPP dataset comprises diverse protein types, each with varying micrograph sizes. Image size ranges from as low as (3710, 3710) to as high as (7676, 7420). For the uniformity in the training process, we resized all the micrographs to (1024, 1024) after denoising them and before feeding them to the U-Net model. From the ground truth coordinate files in the .csv format, containing information like centers of the particles and corresponding diameters, we created a separate ground-truth segmentation mask for each micrograph. This mask was then resized to (1024, 1024). The input micrograph was fed to the network for training while the ground truth segmentation mask was utilized as a target and compared with the output segmentation mask for calculation of loss. **Fig. 7** shows a sample denoised image and its corresponding ground truth segmentation mask.

**Fig. 7.**
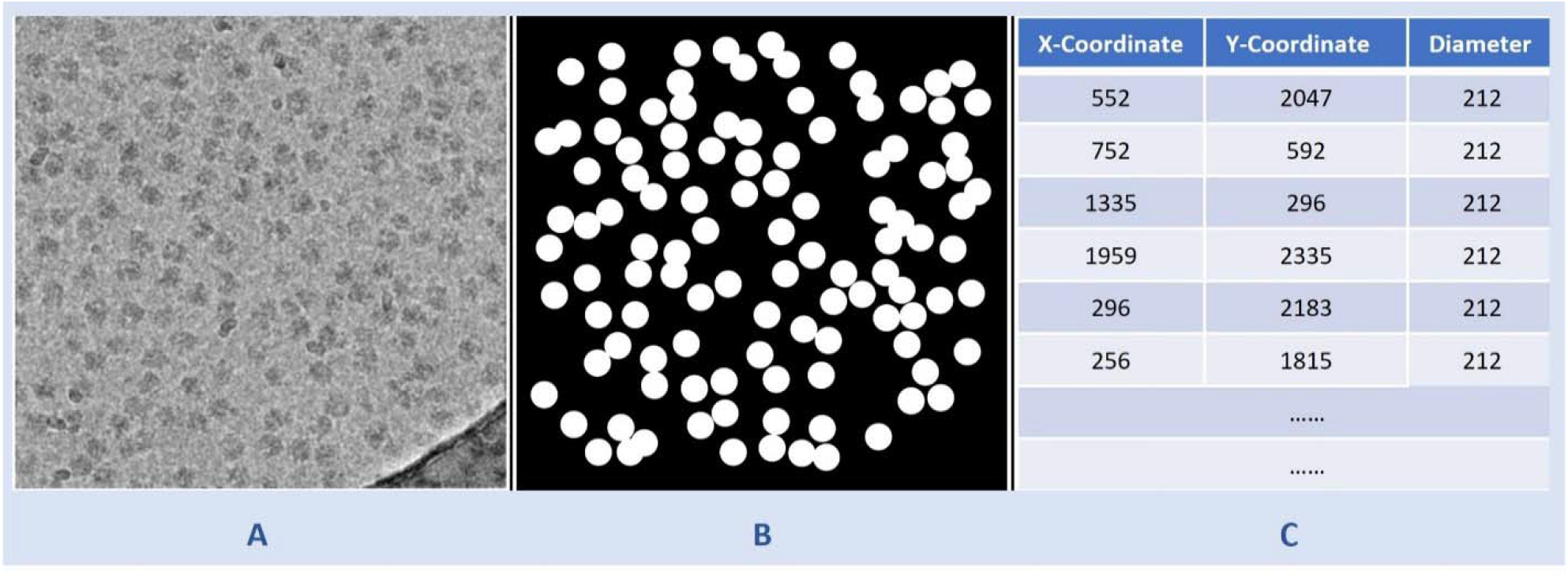
Illustration of data preparation for training the U-Net model. **(A)** A denoised cryo-EM micrograph (EMPIAR ID 10406) as input. **(B)** The ground truth segmentation mask. **(C)** The information from the ground truth coordinate file with x-coordinate and y-coordinate of center of protein particles and corresponding diameters used to generate (B).

### 4. Training

The attention-gated U-Net of CryoSegNet was trained using denoised and resized micrographs of 22 different EMPIAR IDs from CryoPPP dataset. The training was done with a batch size of 6, learning rate of 0.0001 for 200 epochs with a combined loss function of the dice loss and binary cross entropy on NVIDIA A100 80GB GPU.

## Data availability

The dataset for this study is available on https://github.com/BioinfoMachineLearning/cryoppp and https://zenodo.org/record/7934683

## Code availability

The source code is available on https://github.com/jianlin-cheng/CryoSegNet

## Acknowledgements

We thank the entire EMPIAR team for hosting the Cryo-EM data archive. Thanks to the researchers who deposited their cryo-EM images into EMPIAR for public use. We are also thankful to Dr. Filiz Bunyak for her valuable insights on particle picking. Special thanks go to Ali Punjani and his team for developing CryoSPARC, which was extensively used in the validation of results.

## Author contributions

J.C. conceived and conceptualized this research; R.G., A.D., and J.C. designed the deep learning method; J.C and L.W. provided guidance on the experiment and evaluation; R.G. wrote the scripts and codes to preprocess dataset, implemented, trained, and tested the method, and collected the data; R.G., J.C., L.W., and A.D. analyzed the data; R.G. and A.D. drafted the manuscript; J.C. and L.W. revised manuscript. All authors edited the manuscript.

