## Supplementary File for "Accurate cryo-EM protein particle picking by integrating the foundational AI image segmentation model and specialized U-Net"

**This supplementary information contains Supplementary Notes 1-3, Supplementary Figures 1-2 and Supplementary Tables 1-7.**

***Supplementary Note S1:*** ***Parameters in CrYOLO and Topaz***

There are few parameters in CrYOLO and Topaz that affect the quality and number of picked protein particles. While using CrYOLO, two parameters can significantly impact the results. One such parameter is the confidence threshold, which has a default value of 0.3. Increasing this parameter leads to a lower number of picked particles, while a value less than 0.3 tends to result in a larger number of picked particles. We tested various threshold values for each EMPIAR ID in the test dataset, and have utilized the default threshold value that yielded the best results. Another parameter influencing particle picking in CrYOLO is the network architecture it offers. Among the three different architectures, namely “YoLO”, “CrYOLO” and “PhosaurusNet”, we opted for PhosaurusNet, as it can effectively pick small particles and eliminate picking on the carbon edge.

In the case of Topaz, there are also a few parameters that can affect the results of particle picking. One major parameter is the architecture, and we utilized the ResNet16 architecture for particle picking. Another critical parameter is the "radius of extracted regions," which significantly influences the number of picked particles. The default value of 7 often leads to overlapping and duplicates. For instance, in EMPIAR ID 10028 using default value of 7 leads to 169,733 particles. Increasing this value to 10, provides 88,771 particles reducing the duplicates/overlapping particles in high number. Further, increasing the threshold value to 20 leads to very minimal duplicates/overlapping particles yielding only 52,588 particles. For EMPIAR ID 10093 and EMPIAR ID 10017 we used the radius of extracted regions as 15 and for rest of the EMPIAR IDs value of 12 yielded the minimum number of duplicates/overlapping particles. This parameter selection depends from person to person, but the basic idea is to increase the radius parameter by some value and observe if there are any duplicates. The best value can be determined by finding the size and total structure weight of the protein.

***Supplementary Note S2:*** ***Fine-tuning of CrYOLO, Topaz and CryoSegNet***

We conducted fine-tuning on three different methods, namely CrYOLO, Topaz, and CryoSegNet, to assess their performance enhancement on a new dataset. For training and validation, we utilized 20 labeled micrographs per EMPIAR ID from the test dataset. Subsequently, we determined the resolution of the reconstructed density maps for each EMPIAR ID using the three methods. The average resolution remained unchanged for CrYOLO, but it improved from 5.16 Å to 4.85 Å for Topaz and 4.94 Å to 4.42 Å for CryoSegNet when tested on a small set of micrographs. The detailed results are presented in **Supplementary Table S5.** We then applied the fine-tuned models to reconstruct density maps for particles picked on full micrographs. In this scenario, CrYOLO showed an improvement in average resolution from 3.85 Å to 3.82 Å, CryoSegNet improved from 3.32 Å to 3.20 Å, while Topaz exhibited no improvement. **Supplementary Table S6** provides a comprehensive overview of the complete experimental results. Notably, fine-tuning CryoSegNet resulted in a significant improvement in resolution for EMPIAR ID 10093 and EMPIAR ID 10017.

***Supplementary Note S3:*** ***An Ablation Study of CryoSegNet and the Foundational AI Model (SAM)***

We performed a series of experiments to compare CryoSegNet and different ways of using the foundational AI model (i.e., SAM) for protein particle picking from cryo-EM micrographs. We first explored directly applying the pretrained SAM in its original form to the cryo-EM micrographs, which yielded very unsatisfactory results due to the inherent new challenges posed by cryo-EM micrographs (very low contrast and a low SNR) not seen in the data used to trained SAM. Only a few protein particles with distinct contrast and high SNR can usually be segmented by SAM. To address this limitation, we then fine-tuned the SAM’s mask decoder by training it on our dataset for 2000 epochs. We conducted this training by using the weights of three versions of SAM (i.e., ViT-H, ViT-L and ViT-B) as start point, respectively. The three fined tuned SAM generally performed better than the original SAM. Notably, the best segmentation results of the three were achieved by fine-tuning the ViT-H model. Some of the results of segmentation by fine-tuned SAM are shown in **Supplementary Figure S1**. Finally, we explored the approach used by CryoSegNet combining the U-Net model with the SAM’s automatic mask generator, which was performed by feeding the output of the former into the latter to generate the segmentation results. All the three approaches above were applied to the same micrographs denoised by the image processing techniques that enhanced the effectiveness of all the approaches. **Supplementary Figure S2** illustrates the particle segmentation results of the three approaches on one typical example (EMPIAR ID 10028), which clearly demonstrate that combining the U-Net with SAM performs much better than the fine-tuned SAM that in turn substantially outperforms the original SAM. The results show that the U-Net is able to convert original cryo-EM micrographs not well understood by SAM to the segmentation maps that can be handled by it well to improve its performance for particle picking.

Moreover, we tested different components and hyperparameters of the U-Net to assess their contributions. We varied the number of encoder and decoder blocks from 4 to 6 and observed the best performance was achieved with 5 blocks. Additionally, we trained the model with and without using the attention gate before the decoder block. Incorporating the attention gate yielded better segmentation results. Further, we varied the size of the input micrograph from 256x256 to 2048x2048 and the best results were obtained with input size of 1024x1024 in terms of the performance and training and testing speed. Finally, we experimented using dropout at the end of each convolutional block of encoder but found that results without dropout were better.

During the comparison with Topaz in terms of the resolution of reconstructed density maps, we explored the various versions of the Topaz architecture to get its best results. It was observed that the results were superior when using ResNet 16 (64 units) in comparison to ResNet 8 (32 units). Additionally, we conducted the experiments by adjusting the particle threshold parameter, ranging from default value 0 to 2 with an increment of 1. These results are presented in **Supplementary Table S7**. A higher threshold can reduce the number of duplicate and overlapped particles but may also reduce the recall of particle the picking. The results show that increasing the threshold does not improve the resolution of the reconstructed density maps. Therefore, the best result of Topaz achieved at the default threshold of 0 is used to compare with CryoSegNet.

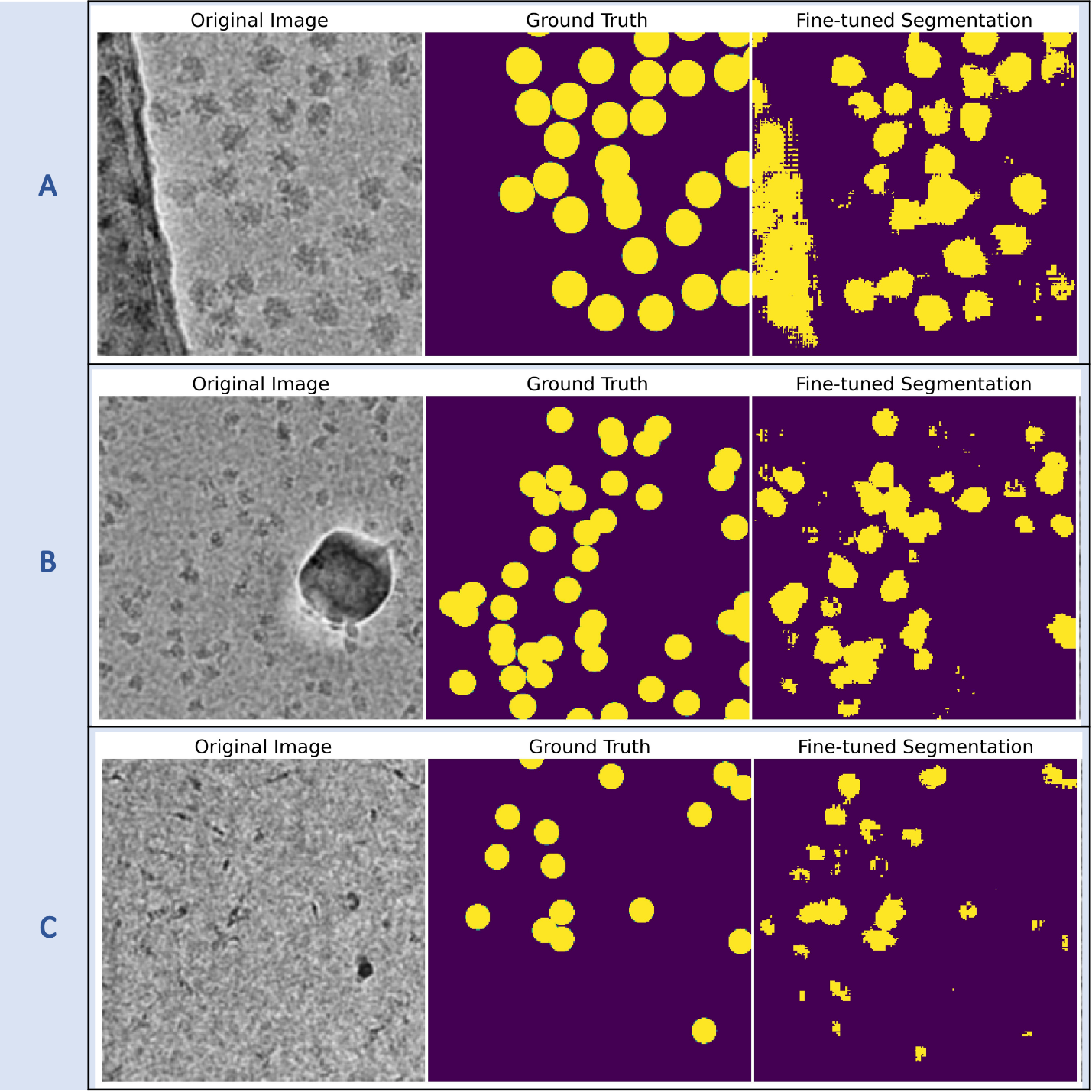

**Supplementary Figure S1**: Segmentation results of fine-tuned SAM **(A)**The whole carbon region is segmented (EMPIAR ID 10532) **(B)** Protein particles of certain orientations are only segmented (EMPIAR ID 10081) **(C)** Some true protein particles are missed and more false positives are segmented (EMPIAR ID 10345)

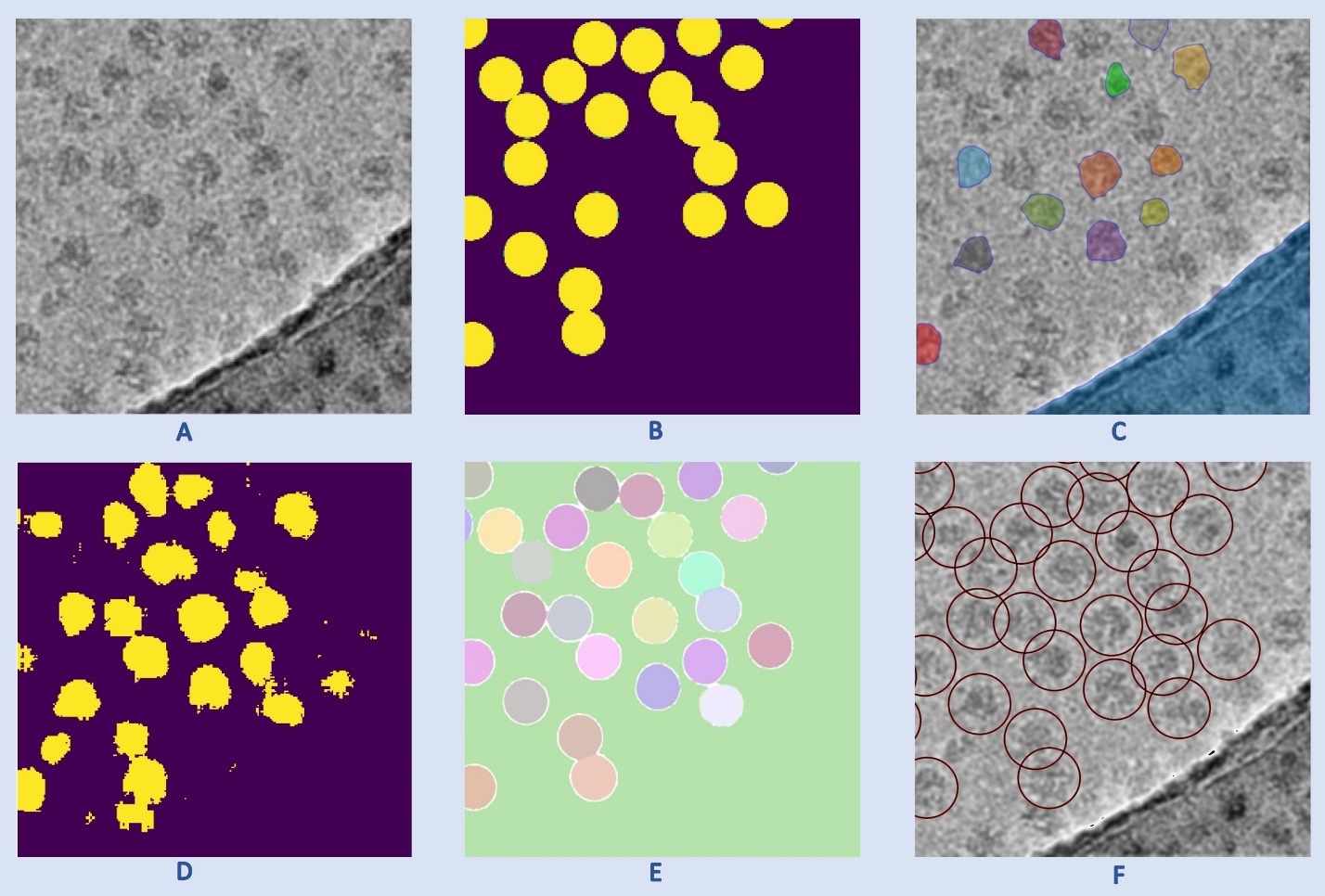

**Supplementary Figure S2:** Results of applying different approaches of using SAM for particle picking. **(A)** Input cryo-EM micrograph (a small patch of a full micrograph from EMPIAR ID 10028). **(B)** Ground truth mask for (A). **(C)** Segmentation result for SAM in its original form. **(D)** Segmentation result for the fine-tuned SAM. **(E)** Segmentation result for SAM used with the U-Net in CryoSegNet. **(F)** Final result for CryoSegNet.

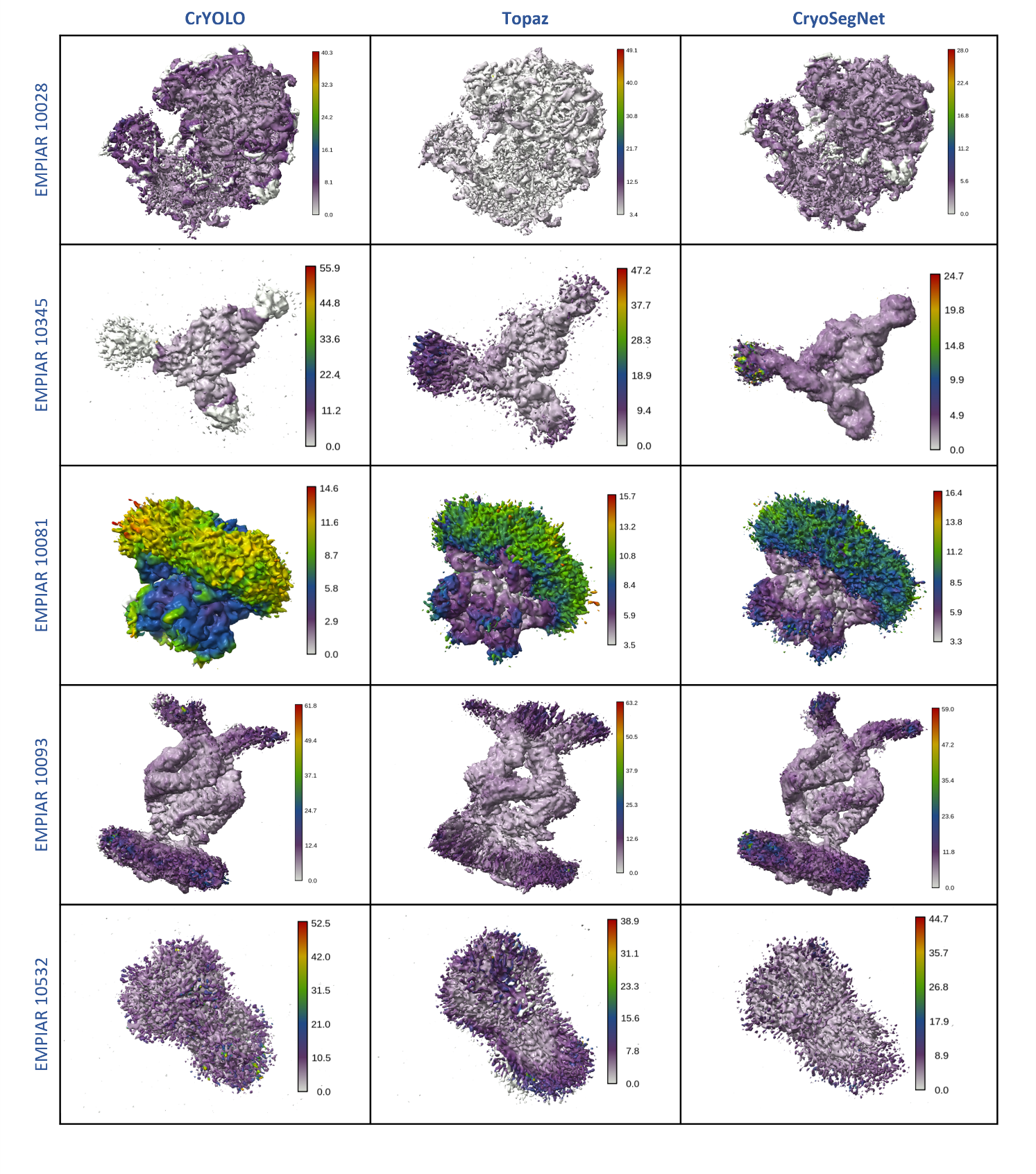

**Supplementary Figure S3:** Evaluation of CrYOLO, Topaz, and CryoSegNet based on the local resolution estimation of 3D density maps. The color scale (in Angstrom) displayed in the right of the map represents high-resolution areas in gray and low-resolution regions in red.

***Supplementary Table S1****: Comparison of precision, recall, F1-score and Dice score for different modules of CryoSegNet*

| **EMPIAR ID** | **Evaluation metrics from output of UNET module** | | | | **Evaluation metrics from output of SAM module** | | | | **Evaluation metrics from Postprocessing module** | | | |
| --- | --- | --- | --- | --- | --- | --- | --- | --- | --- | --- | --- | --- |
|  | **Precision** | **Recall** | **F1 Score** | **Dice Score** | **Precision** | **Recall** | **F1 Score** | **Dice Score** | **Precision** | **Recall** | **F1 Score** | **Dice Score** |
| 10028 | 0.862 | 0.828 | 0.844 | 0.819 | 0.729 | 0.955 | 0.826 | 0.810 | 0.833 | 0.944 | 0.885 | 0.859 |
| 10081 | 0.847 | 0.959 | 0.899 | 0.866 | 0.783 | 0.943 | 0.856 | 0.841 | 0.835 | 0.922 | 0.876 | 0.876 |
| 10345 | 0.569 | 0.872 | 0.688 | 0.610 | 0.656 | 0.911 | 0.763 | 0.725 | 0.746 | 0.920 | 0.824 | 0.743 |
| 11056 | 0.793 | 0.592 | 0.678 | 0.629 | 0.730 | 0.770 | 0.749 | 0.686 | 0.757 | 0.687 | 0.720 | 0.663 |
| 10532 | 0.860 | 0.507 | 0.638 | 0.509 | 0.757 | 0.768 | 0.763 | 0.726 | 0.796 | 0.628 | 0.702 | 0.649 |
| 10093 | 0.454 | 0.944 | 0.613 | 0.570 | 0.607 | 0.661 | 0.633 | 0.592 | 0.716 | 0.515 | 0.600 | 0.537 |
| 10017 | 0.847 | 0.473 | 0.608 | 0.558 | 0.840 | 0.732 | 0.783 | 0.754 | 0.859 | 0.616 | 0.718 | 0.703 |
| Average | 0.747 | 0.739 | 0.710 | 0.652 | 0.729 | 0.820 | 0.768 | 0.733 | 0.792 | 0.747 | 0.761 | 0.719 |

**Supplementary Table S2**: Comparison of precision, recall, F1-score and Dice score for fine-tuned CrYOLO, fine-tuned Topaz and fine-tuned CryoSegNet

| **EMPIAR ID** | **Type of Protein** | **Num. of Labeled Images** | **Num. of Labeled Particles** | **CrYOLO** | | | | **Topaz** | | | | **CryoSegNet** | | | |
| --- | --- | --- | --- | --- | --- | --- | --- | --- | --- | --- | --- | --- | --- | --- | --- |
|  |  |  |  | Precision | Recall | F1 Score | Dice Score | Precision | Recall | F1 Score | Dice Score | Precision | Recall | F1 Score | Dice Score |
| 10028^29^ | Ribosome (80S) | 300 | 26,391 | 0.853 | 0.932 | 0.891 | 0.868 | 0.753 | 0.958 | 0.844 | 0.830 | 0.801 | 0.952 | 0.870 | 0.867 |
| 10081^30^ | Transport | 300 | 39,352 | 0.828 | 0.816 | 0.822 | 0.810 | 0.858 | 0.941 | 0.898 | 0.903 | 0.880 | 0.934 | 0.906 | 0.908 |
| 10345^31^ | Signaling | 295 | 15,894 | 0.874 | 0.880 | 0.877 | 0.789 | 0.639 | 0.904 | 0.749 | 0.728 | 0.756 | 0.926 | 0.833 | 0.770 |
| 11056^32^ | Transport | 305 | 125,908 | 0.776 | 0.540 | 0.637 | 0.571 | 0.793 | 0.707 | 0.748 | 0.676 | 0.849 | 0.800 | 0.824 | 0.796 |
| 10532^33^ | Viral | 300 | 87,933 | 0.736 | 0.838 | 0.784 | 0.752 | 0.812 | 0.704 | 0.754 | 0.738 | 0.821 | 0.719 | 0.766 | 0.694 |
| 10093^34^ | Membrane | 295 | 56,394 | 0.712 | 0.535 | 0.611 | 0.557 | 0.615 | 0.661 | 0.637 | 0.588 | 0.696 | 0.601 | 0.645 | 0.586 |
| 10017^35^ | β-galactosidase | 84 | 49,391 | 0.855 | 0.592 | 0.700 | 0.681 | 0.879 | 0.884 | 0.882 | 0.874 | 0.857 | 0.674 | 0.755 | 0.729 |
| Average | | | | 0.805 | 0.733 | 0.760 | 0.718 | 0.764 | **0.823** | 0.787 | 0.762 | **0.809** | 0.801 | **0.800** | **0.764** |

***Supplementary Table S3****: Comparison of 3D resolution of general CryoSegNet on test dataset with general crYOLO and general Topaz*

| **EMPIAR ID** | **Number of Micrographs** | **Method** | **Without Select 2D** | | | | **With Select 2D** | | | |
| --- | --- | --- | --- | --- | --- | --- | --- | --- | --- | --- |
|  |  |  | **Resolution for 3 Trials (Å)** | | | **Number of Particles** | **Resolution for 3 Trials (Å)** | | | **Number of Particles** |
|  |  |  | **1** | **2** | **3** |  | **1** | **2** | **3** |  |
| 10028 | 300 | *CrYOLO* | **4.13** | 4.14 | 4.13 | 32,687 | 4.12 | **4.11** | 4.11 | 31,699 |
|  |  | *Topaz* | 4.01 | 4.05 | **3.98** | 52,588 | **3.93** | 4.02 | 3.96 | 35,514 |
|  |  | *CryoSegNet* | **2.72** | 2.72 | 2.72 | 47,764 | **2.72** | 2.72 | 2.72 | 45,218 |
| 10081 | 300 | *CrYOLO* | **5.65** | 5.65 | 5.69 | 44,440 | **4.97** | 5.57 | 5.60 | 36,821 |
|  |  | *Topaz* | 6.29 | 6.32 | **6.13** | 58,217 | **5.08** | 5.13 | 5.09 | 37,808 |
|  |  | *CryoSegNet* | **4.58** | 4.59 | 4.62 | 60,158 | **4.16** | 4.17 | 4.20 | 44,819 |
| 10345 | 295 | *CrYOLO* | **3.98** | 3.98 | 4.30 | 15,821 | **3.83** | 3.98 | 4.00 | 11,369 |
|  |  | *Topaz* | 3.89 | 3.84 | **3.73** | 29,208 | **3.64** | 3.68 | 3.65 | 21,343 |
|  |  | *CryoSegNet* | **3.48** | 3.51 | 3.49 | 25,919 | 2.89 | **2.84** | 2.93 | 15,209 |
| 11056 | 305 | *CrYOLO* | 9.22 | 9.19 | **8.98** | 60,648 | 8.65 | **8.32** | 8.65 | 43,599 |
|  |  | *Topaz* | 8.23 | **8.11** | 8.18 | 98,680 | **8.03** | 8.06 | 8.10 | 66,651 |
|  |  | *CryoSegNet* | 7.88 | **7.83** | 7.92 | 71,342 | 7.21 | **7.13** | 7.16 | 53,073 |
| 10532 | 300 | *CrYOLO* | **4.23** | 4.29 | 4.24 | 46,162 | 4.12 | **4.08** | 4.10 | 29,434 |
|  |  | *Topaz* | 4.63 | **4.54** | 4.67 | 73,196 | **4.23** | 4.31 | 4.26 | 38,372 |
|  |  | *CryoSegNet* | 4.13 | 4.16 | **4.09** | 67,219 | 3.95 | **3.89** | 3.93 | 30,155 |
| 10093 | 295 | *CrYOLO* | 7.36 | **7.27** | 7.47 | 43,305 | **6.87** | 6.91 | 7.01 | 33,183 |
|  |  | *Topaz* | **6.35** | 6.51 | 6.42 | 110,577 | **6.12** | 6.18 | 6.14 | 61,698 |
|  |  | *CryoSegNet* | 7.33 | 7.42 | **7.27** | 43,886 | **6.99** | 7.17 | 7.01 | 27,745 |
| 10017 | 84 | *CrYOLO* | 5.14 | **4.99** | 5.11 | 54,263 | 4.87 | **4.84** | 4.89 | 47,704 |
|  |  | *Topaz* | **5.13** | 5.18 | 5.21 | 52,875 | **5.08** | 5.11 | 5.09 | 45,511 |
|  |  | *CryoSegNet* | 6.91 | **6.90** | 6.99 | 11,961 | 6.96 | 6.90 | **6.86** | 10,026 |

***Supplementary Table S4:*** *Comparison of 3D resolution of general CryoSegNet on full set of micrographs with general crYOLO and general Topaz*

| **EMPIAR ID** | **Number of Micrographs** | **Method** | **Without Select 2D** | | | | **With Select 2D** | | | | **Original EMPIAR** | |
| --- | --- | --- | --- | --- | --- | --- | --- | --- | --- | --- | --- | --- |
|  |  |  | **Resolution for 3 Trials (Å)** | | | **Number of Particles** | **Resolution for 3 Trials (Å)** | | | **Number of Particles** | **Resolution (Å)** | **Number of Particles** |
|  |  |  | **1** | **2** | **3** |  | **1** | **2** | **3** |  |  |  |
| 10028 | 600 | *CrYOLO* | 3.99 | 4.00 | **3.97** | 65,376 | 3.99 | **3.94** | 3.96 | 63,562 | 3.20 | 105,247 |
|  |  | *Topaz* | **2.72** | 2.72 | 2.72 | 104,652 | **2.72** | 2.72 | 2.72 | 96,352 |  |  |
|  |  | *CryoSegNet* | **2.72** | 2.72 | 2.72 | 93,881 | **2.72** | 2.72 | 2.72 | 92,532 |  |  |
| 10345 | 1644 | *CrYOLO* | **3.56** | 3.65 | 3.67 | 50,506 | **3.54** | 3.54 | 3.55 | 40,047 | 3.51 | 84,266 |
|  |  | *Topaz* | **3.50** | 3.55 | 3.52 | 102,977 | 3.48 | 3.46 | **3.45** | 87,472 |  |  |
|  |  | *CryoSegNet* | 2.79 | **2.74** | 2.75 | 120,357 | **2.67** | 2.70 | 2.69 | 73,377 |  |  |
| 10081 | 997 | *CrYOLO* | 4.32 | 4.28 | **4.26** | 148,488 | 4.20 | 4.19 | **4.15** | 123,963 | 3.50 | 55,870 |
|  |  | *Topaz* | **4.34** | 4.34 | 4.39 | 171,396 | **4.06** | 4.11 | 4.08 | 130,941 |  |  |
|  |  | *CryoSegNet* | 3.98 | **3.95** | 4.03 | 202,988 | 3.51 | **3.45** | 3.47 | 153,333 |  |  |
| 10532 | 1556 | *CrYOLO* | 3.26 | **3.25** | 3.28 | 232,220 | **3.22** | 3.25 | 3.22 | 161,497 | 2.90 | 128,305 |
|  |  | *Topaz* | 3.63 | 3.61 | **3.52** | 362,115 | 3.23 | **3.22** | 3.23 | 206,460 |  |  |
|  |  | *CryoSegNet* | 3.47 | **3.42** | 3.47 | 181,259 | 3.32 | 3.22 | **3.20** | 90,477 |  |  |
| 10093 | 1873 | *CrYOLO* | 4.61 | **4.54** | 4.58 | 264,447 | **4.41** | 4.42 | 4.46 | 192,337 | 3.55 | 175,314 |
|  |  | *Topaz* | 4.63 | **4.55** | 4.58 | 801,208 | 4.42 | **4.40** | 4.43 | 437,235 |  |  |
|  |  | *CryoSegNet* | 4.90 | **4.70** | 4.76 | 267,983 | 4.58 | **4.54** | 4.61 | 169,330 |  |  |

***Supplementary Table S5:*** *Comparison of 3D resolution of fine-tuned CryoSegNet on test dataset with fine-tuned crYOLO and fine-tuned Topaz*

| **EMPIAR ID** | **Number of Micrographs** | **Method** | **Without Select 2D** | | | | **With Select 2D** | | | |
| --- | --- | --- | --- | --- | --- | --- | --- | --- | --- | --- |
|  |  |  | **Resolution for 3 Trials (Å)** | | | **Number of Particles** | **Resolution for 3 Trials (Å)** | | | **Number of Particles** |
|  |  |  | **1** | **2** | **3** |  | **1** | **2** | **3** |  |
| 10028 | 300 | *CrYOLO* | 4.13 | **4.10** | 4.14 | 31,575 | 4.10 | **4.08** | 4.14 | 31,157 |
|  |  | *Topaz* | **2.72** | 2.72 | 2.72 | 56,431 | **2.72** | 2.72 | 2.72 | 55,868 |
|  |  | *CryoSegNet* | **2.72** | 2.72 | 2.72 | 54,665 | **2.72** | 2.72 | 2.72 | 54,127 |
| 10081 | 300 | *CrYOLO* | 6.26 | 6.33 | **6.18** | 34,543 | 6.14 | **5.87** | 6.12 | 29,682 |
|  |  | *Topaz* | 5.29 | 5.33 | **5.28** | 52,824 | 4.85 | **4.76** | 4.81 | 43,081 |
|  |  | *CryoSegNet* | **4.28** | 4.35 | 4.31 | 53,450 | **4.00** | 4.02 | 4.08 | 45,910 |
| 10345 | 295 | *CrYOLO* | 4.01 | 4.02 | **3.99** | 14,018 | 3.93 | 3.90 | **3.89** | 13,875 |
|  |  | *Topaz* | 3.92 | **3.86** | 3.91 | 37,843 | **3.67** | 3.72 | 3.84 | 23,843 |
|  |  | *CryoSegNet* | **3.01** | 3.13 | 3.07 | 29,804 | 2.85 | 2.89 | **2.81** | 25,799 |
| 11056 | 305 | *CrYOLO* | 8.02 | 8.06 | **8.27** | 71,830 | **7.94** | 7.98 | 7.63 | 50,578 |
|  |  | *Topaz* | 7.98 | **7.91** | 8.03 | 107,732 | 7.72 | **7.63** | 7.78 | 66,804 |
|  |  | *CryoSegNet* | 7.79 | **7.71** | 7.77 | 115,025 | **7.44** | 7.49 | 7.45 | 88,221 |
| 10532 | 300 | *CrYOLO* | 3.98 | 3.97 | **3.93** | 113,479 | 3.93 | **3.91** | 3.92 | 47,094 |
|  |  | *Topaz* | **3.99** | 4.03 | 4.01 | 106,737 | 3.92 | **3.88** | 3.94 | 50,377 |
|  |  | *CryoSegNet* | 3.95 | **3.91** | 3.96 | 99,925 | 3.88 | 3.86 | **3.81** | 45,999 |
| 10093 | 295 | *CrYOLO* | **6.40** | 6.40 | 6.53 | 35,428 | 6.36 | **6.24** | 6.43 | 27,577 |
|  |  | *Topaz* | 6.91 | **6.86** | 6.89 | 53,440 | 6.38 | 6.35 | **6.27** | 41,797 |
|  |  | *CryoSegNet* | **5.46** | 5.55 | 5.51 | 45,801 | 5.01 | **4.98** | 4.92 | 31,023 |
| 10017 | 84 | *CrYOLO* | **5.51** | 5.55 | 5.62 | 31,131 | 5.42 | **5.32** | 5.38 | 28,053 |
|  |  | *Topaz* | 5.16 | **5.09** | 5.13 | 49,720 | **5.02** | 5.03 | 5.08 | 44,972 |
|  |  | *CryoSegNet* | 5.29 | **5.27** | 5.33 | 38,349 | 5.26 | 5.23 | **5.20** | 31,941 |

**Supplementary Table S6:** Comparison of 3D resolution of fine-tuned CryoSegNet on full set of micrographs with fine-tuned crYOLO and fine-tuned Topaz

| **EMPIAR ID** | **Number of Micrographs** | **Method** | **Without Select 2D** | | | | **With Select 2D** | | | | **Original EMPIAR** | |
| --- | --- | --- | --- | --- | --- | --- | --- | --- | --- | --- | --- | --- |
|  |  |  | **Resolution for 3 Trials (Å)** | | | **Number of Particles** | **Resolution for 3 Trials (Å)** | | | **Number of Particles** | **Resolution (Å)** | **Number of Particles** |
|  |  |  | **1** | **2** | **3** |  | **1** | **2** | **3** |  |  |  |
| 10028 | 600 | *CrYOLO* | 3.96 | 3.98 | **3.96** | 62,923 | 3.95 | **3.94** | 3.95 | 62,289 | 3.20 | 105,247 |
|  |  | *Topaz* | **2.72** | 2.72 | 2.72 | 148,955 | **2.72** | 2.72 | 2.72 | 143,074 |  |  |
|  |  | *CryoSegNet* | **2.72** | 2.72 | 2.72 | 109,487 | **2.72** | 2.72 | 2.72 | 107,359 |  |  |
| 10345 | 1644 | *CrYOLO* | 3.59 | 3.98 | **3.56** | 49,526 | 3.58 | 3.81 | **3.55** | 47,893 | 3.51 | 84,266 |
|  |  | *Topaz* | **3.55** | 3.58 | 3.67 | 203,001 | 3.50 | **3.49** | 3.56 | 144,492 |  |  |
|  |  | *CryoSegNet* | **2.78** | 2.82 | 2.79 | 149,961 | 2.73 | **2.67** | 2.69 | 130,750 |  |  |
| 10081 | 997 | *CrYOLO* | 4.54 | 4.42 | **4.40** | 118,455 | 4.42 | **4.27** | 4.32 | 101,165 | 3.50 | 55,870 |
|  |  | *Topaz* | **4.20** | 4.25 | 4.27 | 174,311 | 4.13 | **4.02** | 4.06 | 146,711 |  |  |
|  |  | *CryoSegNet* | **3.75** | 3.77 | 3.81 | 181,953 | 3.53 | 3.58 | **3.47** | 159,577 |  |  |
| 10532 | 1556 | *CrYOLO* | **3.76** | 3.76 | 3.86 | 594,868 | 3.27 | **3.21** | 3.26 | 181,292 | 2.90 | 128,305 |
|  |  | *Topaz* | **3.52** | 3.68 | 3.54 | 583,214 | 3.28 | 3.28 | **3.22** | 238,380 |  |  |
|  |  | *CryoSegNet* | 3.61 | **3.42** | 3.52 | 350,194 | 3.27 | **3.18** | 3.25 | 180,448 |  |  |
| 10093 | 1873 | *CrYOLO* | 4.16 | 4.18 | **4.16** | 223,829 | 4.14 | **4.11** | 4.12 | 173,576 | 3.55 | 175,314 |
|  |  | *Topaz* | **4.48** | 4.50 | 4.51 | 590,917 | **4.40** | 4.46 | 4.43 | 339,233 |  |  |
|  |  | *CryoSegNet* | 4.13 | **4.09** | 4.12 | 278,093 | **3.94** | 3.97 | 3.95 | 220,776 |  |  |

**Supplementary Table S7**: Topaz false positive filtering using different log-likelihood thresholds

| EMPIAR ID | Number of Micrographs | Particle Threshold Parameter 0 | | Particle Threshold Parameter 1 | | Particle Threshold Parameter 2 | |
| --- | --- | --- | --- | --- | --- | --- | --- |
|  |  | Best Resolution (Å) | Number of Particles | Best Resolution (Å) | Number of Particles | Best Resolution (Å) | Number of Particles |
| 10028 | 600 | **2.72** | 273,432 | 2.72 | 165,685 | 2.72 | 108,709 |
| 10345 | 1644 | **3.46** | 245,255 | 3.47 | 170,469 | 3.48 | 83,890 |
| 10081 | 997 | 4.19 | 215,631 | **4.16** | 153,209 | 4.24 | 143,437 |
| 10532 | 1556 | **3.27** | 371,285 | 3.34 | 293,112 | 3.33 | 222,685 |
| 10093 | 1873 | **4.37** | 597,601 | 4.48 | 467,788 | 4.59 | 206,796 |
